# Comprehensive characterization of internal and cuticle surface microbiota of laboratory-reared F_1_ *Anopheles albimanus* originating from different sites

**DOI:** 10.1101/2020.06.02.129619

**Authors:** Nsa Dada, Ana Cristina Benedict, Francisco López, Juan C. Lol, Mili Sheth, Nicole Dzuris, Norma Padilla, Audrey Lenhart

## Abstract

**Background:** Research on mosquito-microbe interactions may lead to new tools for mosquito and mosquito-borne disease control. To date, such research has largely utilized laboratory-reared mosquitoes that typically lack the microbial diversity of wild populations. A logical progression in this area involves working under controlled settings using field-collected mosquitoes or, in most cases, their progeny. Thus, an understanding of how laboratory colonization affects the assemblage of mosquito microbiota would aid in advancing mosquito microbiome studies and their applications beyond laboratory settings.

**Methods:** Using high throughput 16S rRNA amplicon sequencing, we characterized the internal and cuticle surface microbiota of F_1_ progeny of wild-caught adult *Anopheles albimanus* from four locations in Guatemala. A total of 132 late instar larvae and 135 2-5day old, non-blood-fed virgin adult females that were reared under identical laboratory conditions, were pooled (3 individuals/pool) and analyzed.

**Results:** Results showed geographical heterogeneity in both F_1_ larval internal (*p*=0.001; pseudo-*F* = 9.53) and cuticle surface (*p*=0.001; pseudo-*F* = 8.51) microbiota, and only F_1_ adult cuticle surface (*p*=0.001; pseudo-*F* = 4.5) microbiota, with a more homogenous adult internal microbiota (*p*=0.12; pseudo-*F* = 1.6) across collection sites. Overall, ASVs assigned to *Leucobacter, Thorsellia, Chryseobacterium* and uncharacterized *Enterobacteriaceae*, dominated F_1_ larval internal microbiota, while *Acidovorax, Paucibacter*, and uncharacterized *Comamonadaceae*, dominated the larval cuticle surface. F_1_ adults comprised a less diverse microbiota compared to larvae, with ASVs assigned to the genus *Asaia* dominating both internal and cuticle surface microbiota, and constituting at least 70% of taxa in each microbial niche.

**Conclusions:** These results suggest that location-specific heterogeneity in filed mosquito microbiota can be transferred to F_1_ progeny under normal laboratory conditions, but this may not last beyond the F_1_ larval stage without adjustments to maintain field-derived microbiota. Our findings provide the first comprehensive characterization of laboratory-colonized F_1_ *An. albimanus* progeny from field-derived mothers. This provides a background for studying how parentage and environmental conditions differentially or concomitantly affect mosquito microbiome composition, and how this can be exploited in advancing mosquito microbiome studies and their applications beyond laboratory settings.

## Background

Mosquitoes contain microbes that inhabit various tissues, such as the alimentary canal, reproductive organs, and cuticle surface [1]. These microbes are thought to be principally obtained from the mosquito habitat during larval development, and from food sources at the adult stage [1]. In addition to acquisition from larval habitats and/or adult food sources, transovarial bacterial transmission from adult females to their eggs, and transstadial transmission across different immature stages, and into the adult stage, have been demonstrated [2, 3]. As a key component of the mosquito microbiota, environmentally-acquired microbes have been shown to affect mosquito life history traits such as the rate of pupation and adult body size [4]. The mosquito microbiota has also been shown to affect the following aspects of mosquito biology: immunity to human pathogens [5], reproduction [6], insecticide resistance [7, 8], and ultimately vector competence—the mosquito’s ability to acquire, maintain and transmit pathogens [5]. These effects of the microbiota on mosquito biology are being leveraged to develop novel approaches for mosquito-borne disease control [9].

The use of next generation molecular biology tools has resulted in extensive characterization of mosquito microbiota, with the initial focus on bacterial and archaeal components now expanding to eukaryotic microbes [10, 11] and viruses [12, 13]. These advances in mosquito microbiota research have led to field applications of mosquito symbionts for mosquito control. For *Aedes aegypti*, the principal vector of dengue, Zika, chikungunya and yellow fever viruses, mosquito-derived symbionts are now being used to suppress mosquito populations [14] and also being considered to control the spread of pathogens [15]. However, studies exploring mosquito symbionts for malaria control have largely remained at the laboratory stage [16, 17]. Similarly, the microbiota of mosquito vectors in some geographical regions are well characterized and studied compared to those from other regions. In malaria vectors for example, studies on the microbiota have largely focused on Sub-Saharan African species—in particular, *Anopheles gambiae—*and to a lesser extent on those from Southeast Asia [18]. In contrast, the microbiota of Latin American malaria vectors have only recently been comprehensively characterized [7, 8, 19-21], with these studies describing associations between *An. albimanus* microbiota and insecticide resistance [7, 8], and the factors that shape the composition of *An. darlingi, An. albimanus, An. nunetzovari, An. rangeli*, and *An. triannulatus* microbiota [19-21].

To exploit the mosquito microbiota for malaria and malaria vector control, research must successfully advance from laboratory to field settings, a transition which can be fraught with challenges. For example, some malaria vectors such as *An. darlingi, An. vestitipennis*, and *An. gambiae* breed in sites that are small, temporary and often difficult to find and/or access [22-26], making it hard to obtain sufficient immature field mosquitoes for experiments. Where larval habitats are plentiful and easy to find and/or access, the subsequent rearing of field-collected mosquitoes to obtain uniform characteristics can pose additional challenges [27, 28]. Additionally, some malaria vectors belong to species complexes whose members are morphologically indistinguishable [29-32], constituting another layer of complexity that needs to be considered in elucidating mosquito-microbe interactions in malaria vectors.

These challenges, which are common to research on mosquito ecology and control, are often not reported or discussed in mosquito microbiome studies. Several failed attempts at collecting and rearing sufficient immature mosquitoes from the field for our previous study on the role of mosquito microbiota in insecticide resistance resulted in ultimately using either wild-caught adults [7] or F_1_ progeny derived from field-collected adult mosquitoes [8, 33]. While field-caught adult mosquitoes or their F_1_ progeny may offer insights into mosquito-microbe interactions in field scenarios, obtaining adult field-collected mosquitoes with uniform and/or controlled physiological characteristics is usually not feasible. Although geographically associated heterogeneity in microbiota of field-collected mosquitoes has been previously described [34, 35], there is limited information on the fate of field-acquired microbiota after laboratory colonization of field-collected mosquitoes. So far, it has been observed that upon eclosion, newly-emerged laboratory-reared adult mosquitoes show a reduction in bacterial diversity in contrast to earlier developmental stages [36]. In addition, a recent study on the fate of field-acquired microbiota in laboratory-colonized *An. gambiae s*.*l*. showed a reduction in bacterial diversity of the F_5_ progeny (the first point of measurement) that were reared in dechlorinated tap water in contrast to F_0_ [37]. Another study of *Ae. aegypti* from different geographical locations showed no associations between geographical location and microbiota composition after several generations of laboratory colonization [38]. At what point the microbiota of laboratory-colonized mosquitoes become homogenous and whether the microbiota of F_1_ laboratory progeny represent their parental origin—and thus could be used for symbiont-based translational studies—remains largely undescribed.

Here, we present a comprehensive characterization of the microbiota of laboratory-reared F_1_ progeny from field-caught adult *An. albimanus*. These represent unanticipated findings from a larger study that was aimed at characterizing the effects of insecticide exposure and resistance on *An. albimanus* microbiota [8]. We expected little or no immediate loss of geographical heterogeneity in microbial composition upon initial laboratory colonization based on evidence of this type of heterogeneity in field-derived populations [34, 35]. However, our data showed geographical heterogeneity in both F_1_ larval internal and cuticle surface microbiota, and only F_1_ adult cuticle surface microbiota, with a more homogenous adult internal microbiota. These findings lay the foundations for studying how parentage and environmental conditions differentially or concomitantly affect mosquito microbiome composition, and how this can be exploited in advancing mosquito microbiome studies and their applications beyond laboratory settings.

## Results

Using high throughput deep sequencing of the universal bacterial and archaeal 16S rRNA gene, we characterized the microbiota of the internal and cuticle surface microbial niches of laboratory reared *An. albimanus* F_1_ larvae (n=132) and adult (n=135) progeny originating from mothers that were collected from four locations in south central Guatemala (Fig 1). Mosquitoes were processed as pools (3 individuals per pool), resulting in a total of 44 larval and 45 adult pools. F_1_ larvae were from El Tererro, Las Cruces 3 and Las Cruces 4, while F_1_ adults were from Las Cruces 1, Las Cruces 3 and Las Cruces 4 (Table 1). Following quality control of the resulting sequencing reads, amplicon sequence variants (ASVs) were identified and used in downstream analysis. Since adult mosquito microbiota is distinct from that of immature stages [39, 40], ASVs from larvae and adults were analyzed separately, as were ASVs from each microbial niche (internal or cuticle surface).

**Figure 1:**
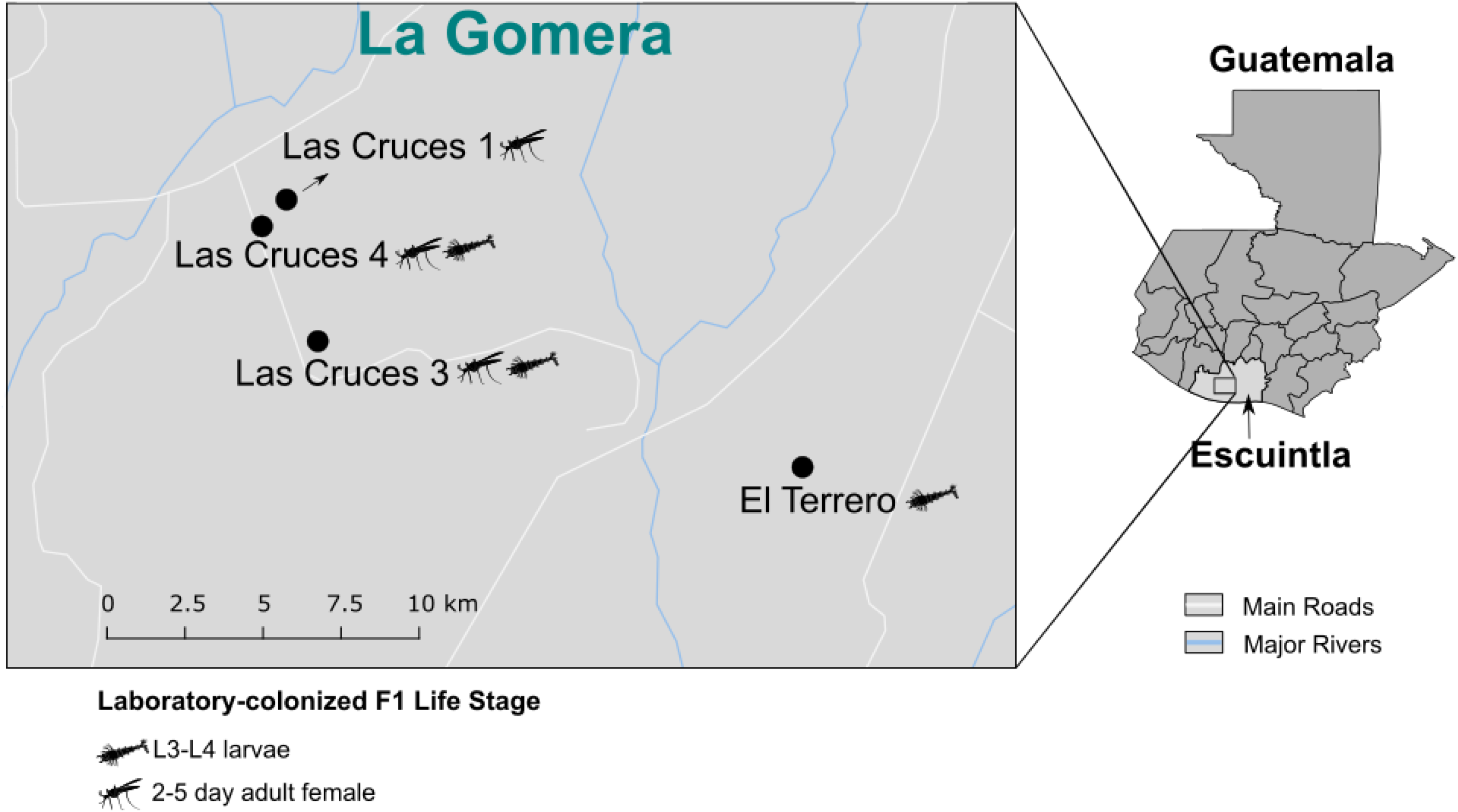
Map showing the geographical origins of F_1_ *Anopheles albimanus*. F_1_ laboratory-colonized mosquitoes were derived from gravid and/or blood fed females collected from each location. Mosquito icons show the geographical origin of the F_1_ life stages studied.

**Table 1.** Descriptive statistics of laboratory colonized F_1_ *An. albimanus* processed, per life stage and collection site.

| Collection site | Number of mosquitoes (number of pools) |  |
| --- | --- | --- |
|  | L3-L4 larvae | Adult ♀s |
| El Terrero | 45 (15) | - |
| Las Cruces 1 | - | 27 (9) |
| Las Cruces 3 | 42 (14) | 45 (15) |
| Las Cruces 4 | 45 (15) | 63 (21) |

### Internal and cuticle surface microbiota of laboratory colonized F_1_ *An. albimanus* larvae differed by geographic origin

Non-pairwise Bray-Curtis distance comparison showed significant differences in internal (p=0.001) and cuticle surface (p=0.001) microbiota between F_1_ larvae from different collection sites. Thus, irrespective of microbial niche, the microbial community structure (composition and relative abundance of ASVs) of F_1_ laboratory-colonized larvae differed by collection location. Pairwise PERMANOVA comparison of Bray-Curtis distances further showed significant differences in microbial community structure in larval internal (q<0.01) and cuticle surface microbiota (q<0.01) between every pair of collection site (Table 2a). This location-driven heterogeneity in microbial community structure was further demonstrated by principal coordinate analysis (PCoA), where F_1_ larval internal and cuticle surface microbiota clustered distinctly by collection site (Fig. 2).

**Table 2.** Pairwise alpha and beta diversity comparisons of laboratory colonized F_1_ *An. albimanus* microbiota from different collection sites. **a**. Pairwise beta (Bray-Curtis) diversity comparison showed significant differences in larval internal and cuticle surface microbiota between collection sites. In contrast, only adult cuticle surface but not internal microbiota were significantly different across collection sites. **b**. Pairwise alpha (Shannon) diversity comparison showed significant differences in larval internal but not cuticle surface microbiota between collection sites (two of the three pairs). In contrast, there was no significant difference in adult internal or cuticle surface microbiota between collection sites. Pairwise alpha and beta diversity comparisons were conducted using Kruskal-Wallis and PERMANOVA (999 permutations) tests respectively, with Benjamini-Hochberg FDR correction (q-value). Significance was determined at q < 0.05. n = No. of pools processed, and each pool comprised three individual mosquitoes.

**a.**
|  | Group 1 | Group 2 | Internal |  |  | Cuticle surface |  |  |
| --- | --- | --- | --- | --- | --- | --- | --- | --- |
|  |  |  | pseudo-F | p-value | q-value | pseudo-F | p-value | q-value |
| Adults | LasCruces1 (n=9) | LasCruces3 (n=15) | 2.21 | 0.088 | 0.161 | **5.80** | **0.001** | **0.003** |
|  | LasCruces1 (n=9) | LasCruces4 (n=21) | 1.92 | 0.107 | 0.161 | **4.97** | **0.004** | **0.005** |
|  | LasCruces3 (n=15) | LasCruces4 (n=21) | 0.85 | 0.443 | 0.443 | **3.07** | **0.005** | **0.005** |
| Larvae | EITerrero<sup>a</sup> (n=15) | LasCruces3 (n=14) | **9.81** | **0.001** | **0.001** | **4.48** | **0.001** | **0.001** |
|  | EITerrero<sup>a</sup> (n=15) | LasCruces4 (n=15) | **7.98** | **0.001** | **0.001** | **11.43** | **0.001** | **0.001** |
|  | LasCruces3 (n=14) | LasCruces4 (n=15) | **10.88** | **0.001** | **0.001** | **11.37** | **0.001** | **0.001** |

<sup>a</sup>Two pools were excluded from the analysis of cuticle surface microbiota following rarefaction

**b.**
|  | Group 1 | Group 2 | Internal |  |  | Cuticle surface |  |  |
| --- | --- | --- | --- | --- | --- | --- | --- | --- |
|  |  |  | H | p-value | q-value | H | p-value | q-value |
| Adults | LasCruces1 (n=9) | LasCruces3 (n=15) | 0.00 | 0.98 | 0.98 | 1.09 | 0.30 | 0.41 |
|  | LasCruces1 (n=9) | LasCruces4 (n=21) | 0.49 | 0.48 | 0.72 | 1.23 | 0.27 | 0.41 |
|  | LasCruces3 (n=15) | LasCruces4 (n=21) | 1.77 | 0.18 | 0.55 | 0.67 | 0.41 | 0.41 |
| Larvae | EITerrero (n=15) | LasCruces3 (n=14) | **6.63** | **0.01** | **0.02** | 4.45 | 0.03 | 0.10 |
|  | EITerrero (n=15) | LasCruces4 (n=15) | **6.94** | **0.01** | **0.02** | 1.87 | 0.17 | 0.26 |
|  | LasCruces3 (n=14) | LasCruces4 (n=15) | 0.49 | 0.48 | 0.48 | 1.10 | 0.29 | 0.29 |

**Figure 2.**
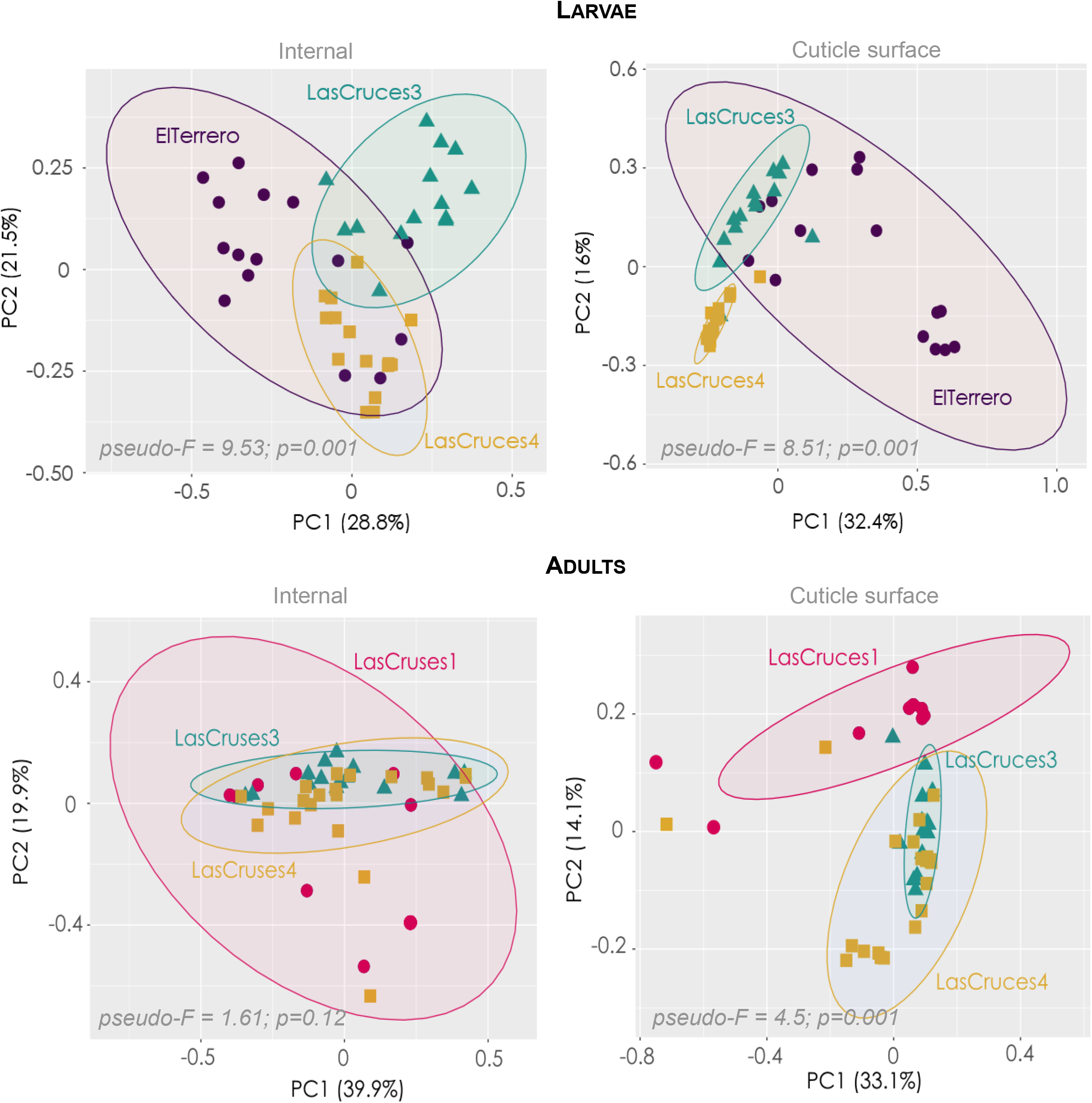
Principal coordinate analysis (PCoA) ordinations of internal and cuticle surface microbiota from F_1_ laboratory-colonized *An. albimanus*. The PCoA plots are based on Bray-Curtis distances between the microbiota of mosquitoes with differing collection sites. Each point on the plot represents the microbial composition of a pool of three individuals, and mosquito pools are color-coded by their origin. For larvae, the first two principal component (PC) axes captured 50% (internal) and 48% (cuticle surface) of the variance in the data, with both internal and cuticle surface microbiota clustering distinctly by collection site. For adults, the first two PC axes captured 59% (internal) and 47% (cuticle surface) of the variance in the data, with cuticle surface but not internal microbiota clustering distinctly by collection site. PERMANOVA statistics are presented at the bottom of each plot.

Non-pairwise Shannon diversity comparisons showed significant differences in internal (p=0.009) but not cuticle surface (p=0.09) microbiota of F_1_ laboratory-colonized larvae from different collection sites, indicating that there was inter-sample variation in the diversity of internal but not cuticle surface microbiota of larvae when all collection sites were taken into consideration. A pairwise Kruskal-Wallis comparison of Shannon diversity indices showed that the inter-sample variation in diversity of larval internal microbiota held true when every pair of collection sites was considered except between Las Cruces 3 and 4 (Table 2b and Suppl. 1). Larvae originating from Las Cruces 3 had the highest internal microbiota diversity, followed by Las Cruces 4 and El Terrero (Suppl. 1)

### Cuticle surface, but not internal, microbiota of laboratory colonized F_1_ adult *An. albimanus* differed by geographic location

Non-pairwise Bray-Curtis diversity comparisons showed significant differences in cuticle surface (p=0.001), but not internal microbiota (p=0.12) between adult F_1_ mosquitoes from different collection sites, suggesting a loss of location-driven heterogeneity in microbial community structure in internal but not cuticle surface microbial niche of laboratory-colonized F_1_ adults. Pairwise PERMANOVA comparisons of Bray-Curtis distances also showed significant differences in microbial community structure of F_1_ adult cuticle surface microbiota (q<0.01) between every pair of collection sites (Table 2a). These results were corroborated by PCoA which showed that F_1_ adult cuticle surface microbiota, but not internal microbiota, clustered distinctly by collection site (Fig. 2).

Non-pairwise Shannon diversity comparisons showed no differences in the internal (p=0.42) or cuticle surface (p=0.4) microbiota of F_1_ adults from different collection sites, indicating that there was little or no inter-sample variation in diversity of F_1_ adult microbiota when all collection sites were taken into consideration. Pairwise Kruskal-Wallis comparisons of Shannon diversity indices also detected no inter-sample variation in diversity of F_1_ adult cuticle surface or internal microbiota when every pair of collection site was considered (Table 2b and Suppl. 1).

### Laboratory-colonized F_1_ *An. albimanus* larvae comprised a rich and diverse microbiota that differed by geographic location

Overall, ASVs from larval internal microbiota were assigned to 180 bacterial taxa, and cuticle surface microbiota to 194 bacterial taxa (suppl 2.). A majority of these taxa across all locations (ranging from 118-139 taxa) were shared between the internal and cuticle surface microbiota (Fig 3a), as well as across collection sites (n=110 for cuticle surface and n=117 for internal microbiota) (Fig 3b). While a majority of the identified microbial taxa were shared between both microbial niches, their abundance was generally higher in internal (Fig 4a) compared to cuticle surface (Fig 4b) microbiota. Although a majority of identified microbial taxa in both internal and cuticle surface microbiota were shared across all locations, their abundance differed by location (Fig 4a, 4b and 5).

**Figure 3.**
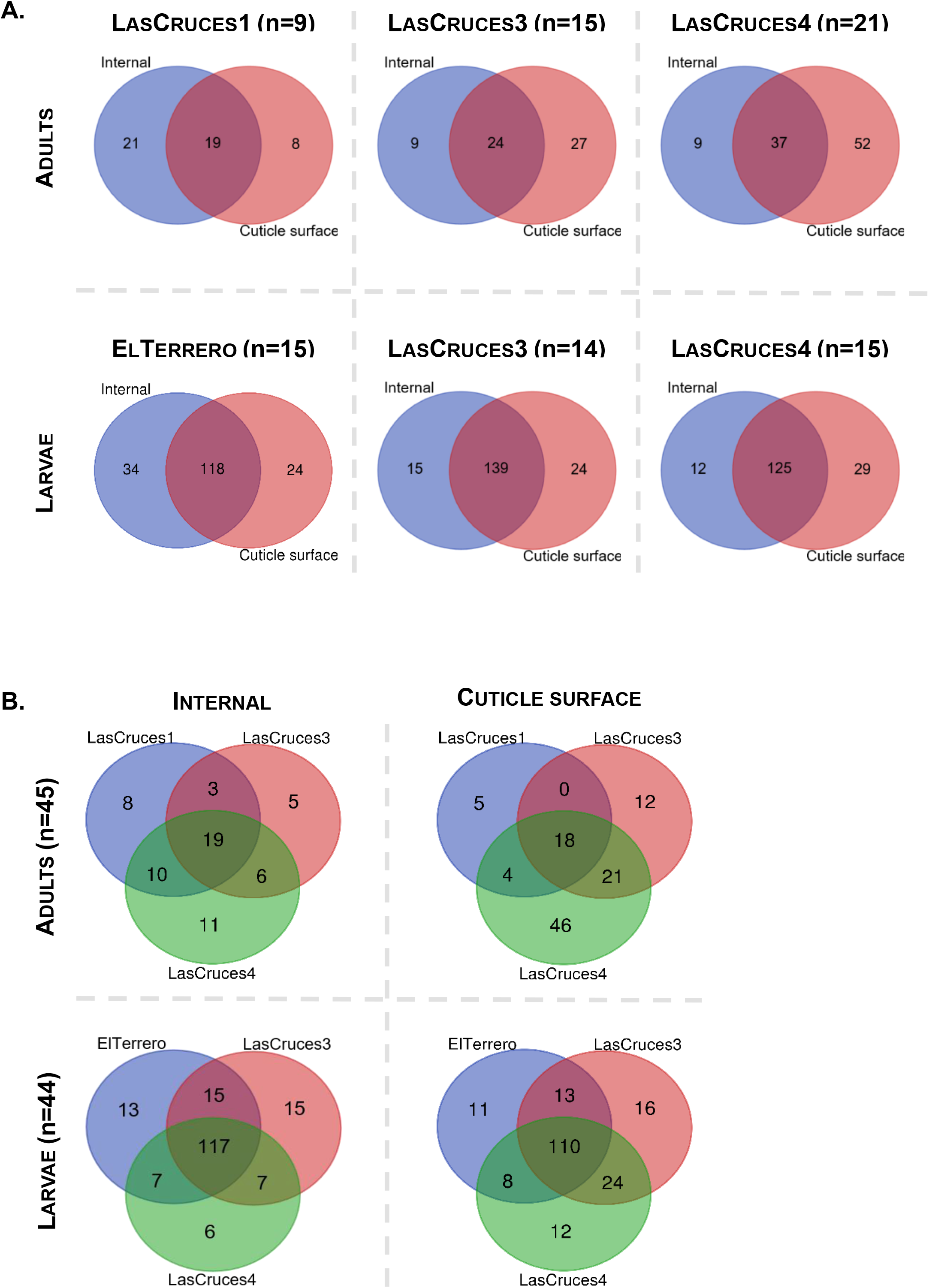
Number of unique and shared microbial taxa between microbial niches (A); number of unique and shared microbial taxa between collection sites (B). The number of taxa shown in the Venn diagram represent bacterial taxa except in the cuticle surface microbiota of adults from Las Cruces 4, where two archaeal taxa were present. n = pools of mosquito samples analyzed per location or microbial.

**Figure 4.**
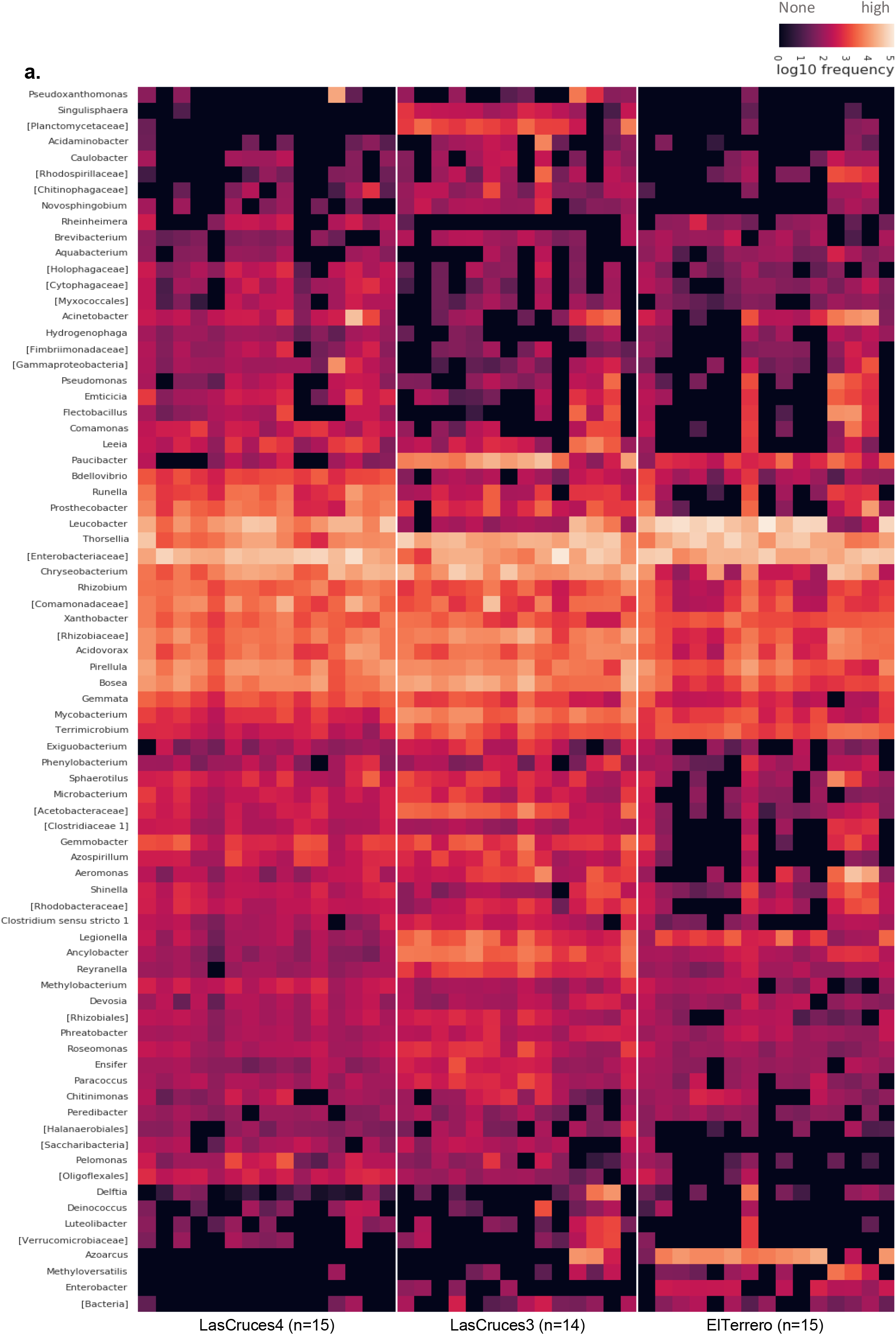
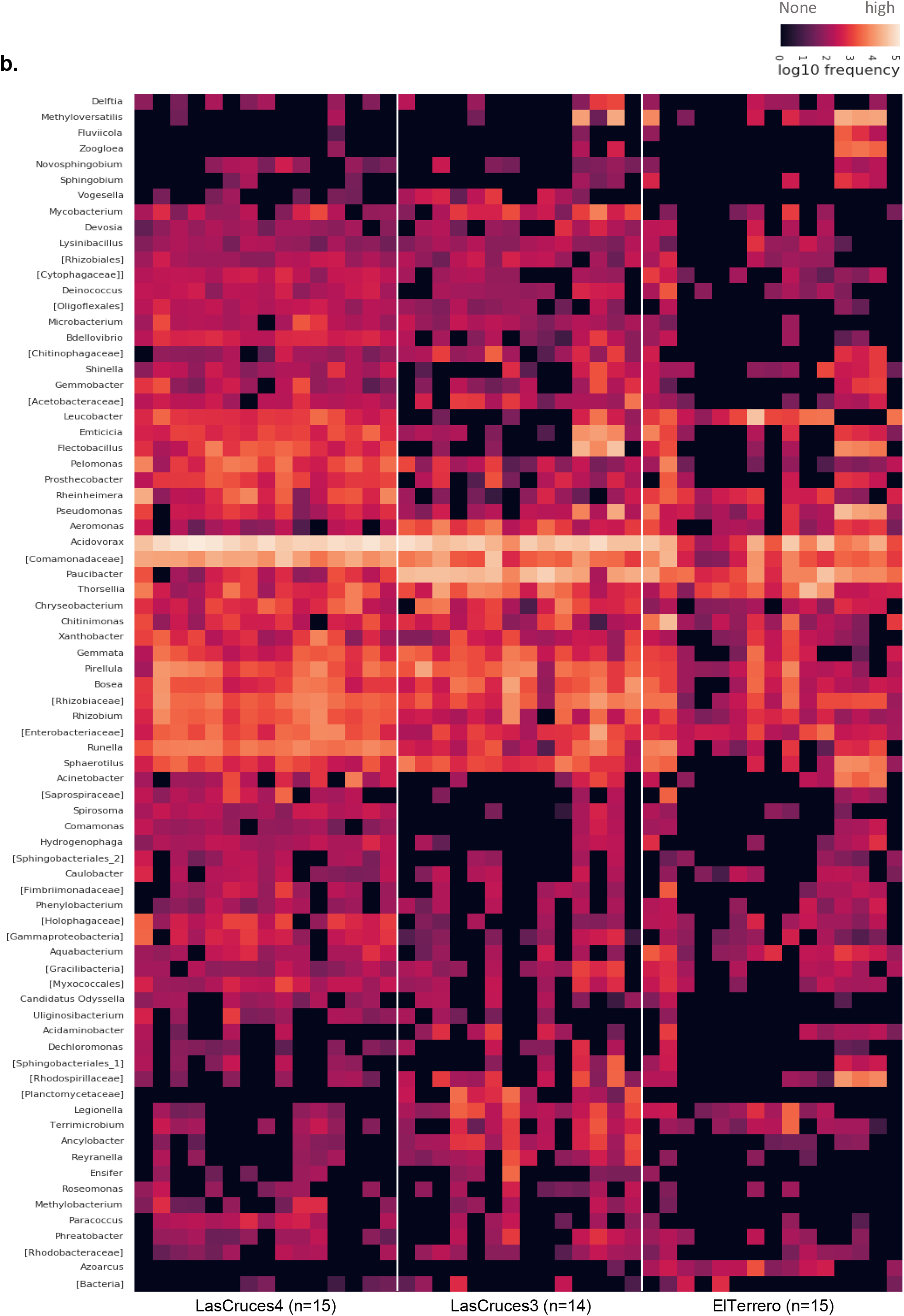
Frequency of ASVs from the internal (a) and cuticle surface (b) microbiota of laboratory-colonized *An. albimanus* F_1_ larvae originating from different collection sites. ASVs were annotated to the genus level or the lowest possible taxonomic level (in square brackets) and are clustered by the average nearest-neighbors chain algorithm. Only taxonomically annotated ASVs with frequencies ≥ 2000 are presented.

**Figure 5.**
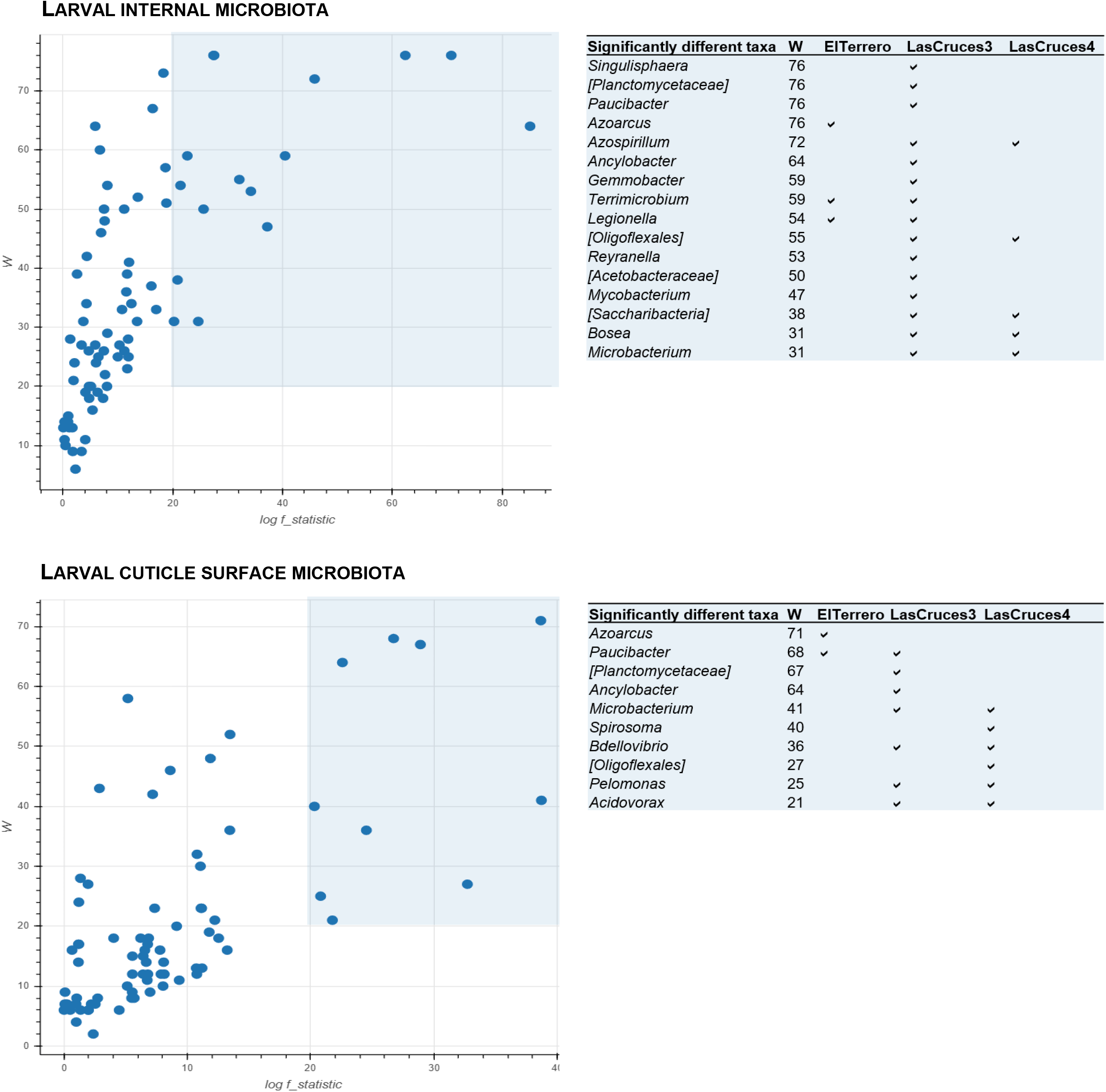
Volcano plots of differentially abundant bacterial taxa in F_1_ laboratory colonized *An. albimanus* larvae originating from three different collection sites. The plots show results of analysis of composition of microbiomes (ANCOM) tests for differentially abundant microbial taxa between collection site, with an effect size set to log F≥20, and a cut-off of differential abundance set to W≥20 (i.e., a taxon was differentially abundant across collection sites if the ratio of its abundance to those of at least 20 other taxa (25% of all included taxa) differed significantly across sites). Differentially abundant taxa are highlighted (blue shaded area) and the taxa names and locations in which they were most abundant are presented in the adjoining tables.

In general, larval internal microbiota was dominated by ASVs identifed as an uncharacterized *Enterobacteriaceae, Leucobacter, Thorsellia*, and *Chryseobacterium* (Fig 4a), together making up over 50% of ASVs (Suppl. 2). In contrast, *Acidovorax*, unchracterized *Comamonadaceae*, and *Paucibacter* (Fig 4b) made up over 50% of ASVs detected on the larval cuticle surface (Suppl. 2).

A few predominant bacterial taxa were present in the larval internal microbial niche across all three collection sites: unclassified *Enterobacteriaceae, Thorsellia, Rhizobium, Xantobacter, Acidovorax* and *Pirellula* (Fig 4a.), with remaining taxa showing different patterns of abundance between collection sites (Fig 5 and Suppl. 2). For example, ASVs assigned to the genus *Azoracus* were predominant in larvae from El Terrero, while *Singulisphaera, Paucibacter, Ancylobacter, Gemmobacter*, and *Rayranella* were predominant in those from Las Cruces 3. Predominant in larvae from Las Cruces 3 and Las Cruces 4 were *Azospirillum, Bosea* and *Microbacterium*; and in those from El Terrero and Las Cruces 3 were *Terrimicrobium* and *Legionella* (Fig 5). Although outside of the cut off limit set for differential abundance, ASVs assigned to the bacterial genera *Leucobacter* were predominant in the internal microbiota of larvae from Las Cruces 4 and El Terrero, but predominant in only three of the 14 sample replicates from Las Cruces 3 (Fig 4a). Similarly, *Chryseobacterium* was predominant in Las Cruces 4 and 3, but only predominant in six of the 16 pools of larvae from El Terrero. Bacterial taxa that were unique to each location comprised <8% of all taxa in larval internal microbiota (Suppl. 3b), and were below the threshold for inclusion in the heatmap and differential abundance testing (Suppl. 2).

Unlike the internal microbial niche, no microbial taxa was predominant in larval cuticle surface microbiota across all three collection sites. However, some taxa showed notable patterns of abundance between locations (Fig 5). These included the genus *Azoarcus*, which was detected at low to moderate frequencies in 13 of 15 pools of larvae from El Terrero, at low frequency in a single pool of larvae from Las Cruces 3, and was not detected at all in Las Cruces 4 (Fig 4b and 6). Similarly, ASVs assigned to the genus *Spirosoma* were detected at moderate frequencies in all pools of larvae from Las Cruces 4, but only in a few pools from the other two locations. ASVs assigned to the genus *Paucibacter* were present at relatively higher abundance in larvae from both Las Cruces 3 and El Terrero compared to those from Las Cruces 4. Those assigned to the genus *Acidovorax* were predominant in larvae from Las Cruces 3 and Las Cruces 4 in contrast to El Terrero. ASVs assigned to *Microbacterium, Bdellovibrio* and *Pelomonas* were present at moderate frequencies in larvae from both Las Cruces 3 and Las Cruces 4 but were not detected in El Terrero(Fig. 5). Bacterial taxa that were unique to each collection site comprised <8% of larval cuticle surface microbiota (Fig 3b), and were below the threshold for inclusion in the heatmap and differential abundance testing (Suppl. 2).

### Laboratory-colonized adult F_1_ *An. albimanus* were comprised of sparse internal and cuticle surface microbiota that were dominated by ASVs assigned to the genus *Asaia*

ASVs from adult internal microbiota were assigned to 62 microbial taxa and cuticle surface microbiota were assigned to 106 microbial taxa. Two of these ASVs which were only present in the cuticle surface microbiota were classified as archaea, while all other remaining ASVs were classified as bacteria (Suppl. 2). Unlike larval microbiota, less than half of the assigned taxa across all locations (ranging from 19-37 taxa) were shared between internal and cuticle surface microbiota (Fig 3a), and only 18 taxa on the cuticle surface and 19 internal taxa were shared across all maternal collection sites (Fig 3b).

Overall, ASVs assigned to the bacterial genus *Asaia* dominated both adult internal and cuticle surface microbiota (Fig 6), constituting at least 70% of taxa in each microbial niche (Suppl. 2). A majority of identified taxa in adult internal, but not cuticle surface, microbiota was detected across all three collection sites, with a few of these taxa present in high abundance across all collection sites (Fig. 6). Across all three collection sites, ASVs assigned to the genera *Acinetobacter, Gluconobacter, Pantoea* and *Pseudomonas* were present in moderate to high abundance in adult internal microbiota in addition to *Asaia* (Fig 6).

**Figure 6.**
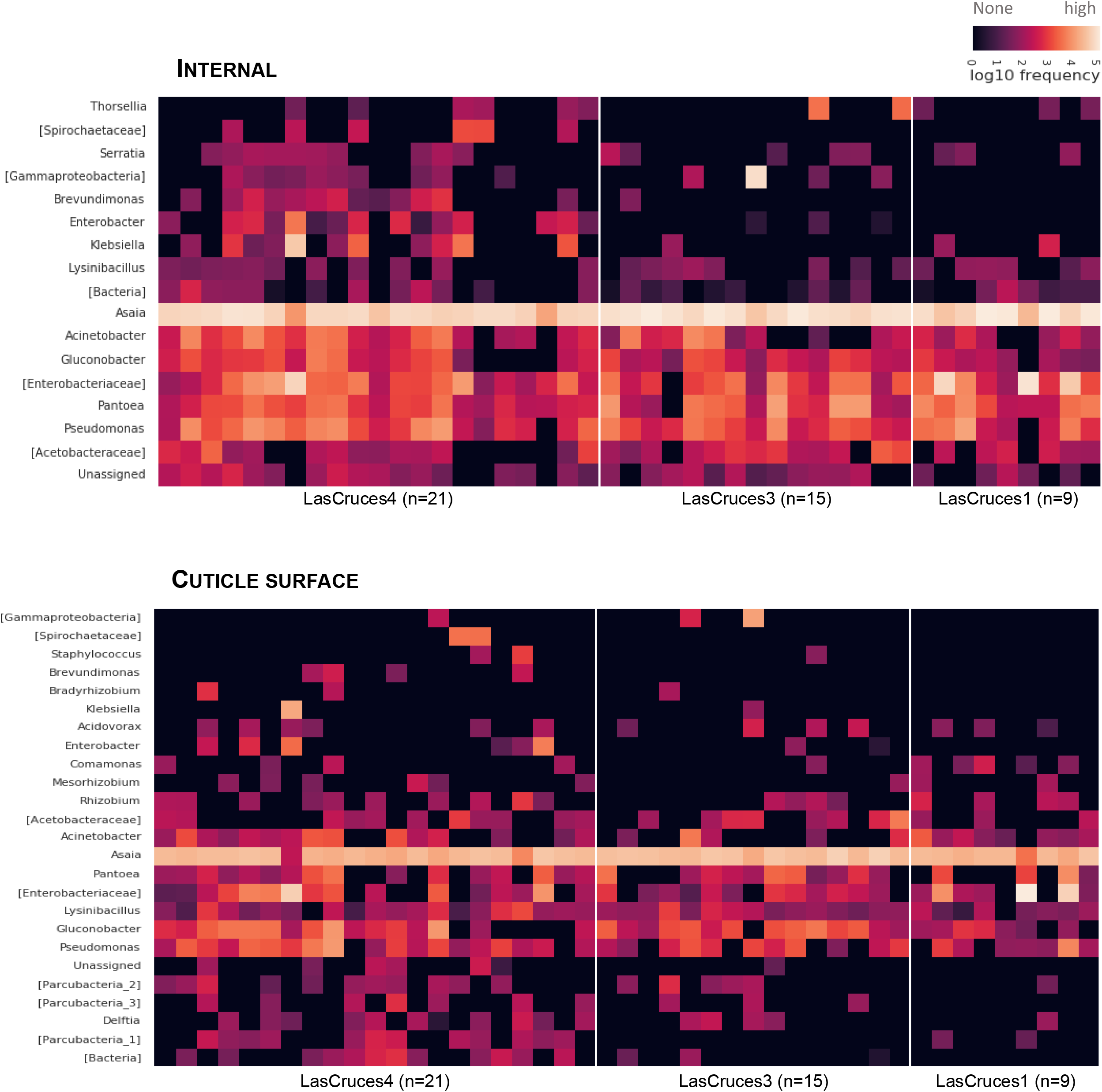
Frequency of ASVs from the internal and cuticle surface microbiota of laboratory-colonized F_1_ adult *An. albimanus* originating from different locations. ASVs were annotated to the genus level or the lowest possible taxonomic level (in square brackets) and are clustered by the average nearest-neighbors chain algorithm. Only taxonomically annotated ASVs with frequencies ≥1000 are presented.

No site-specific microbial taxa were identified in adult cuticle surface microbiota, as the ASV distribution did not meet the criteria for this type of analysis. This was compounded by dominance (>70%) of ASVs that were assigned to the bacterial genus *Asaia* (Suppl. 2). In addition, the cuticle surface microbiota of adults originating from Las Cruces 4 comprised 43% of all adult cuticle surface microbial taxa (Fig 3b), although a majority were of low abundance.

## Discussion

The scientific community is increasingly investigating the role of mosquito microbiota in fighting mosquito-borne diseases [9]. The successful transition of mosquito microbiome research from laboratory to field requires a comprehensive understanding of the dynamics underlying the composition of the microbiota of field-collected mosquitoes and their progeny. We provide a comprehensive characterization of the internal and cuticle surface microbiota of laboratory-reared F_1_ progeny from field-caught adult *An. albimanus* that were collected from different locations. Our results show that while location-driven heterogeneity in the microbial community structure of both internal and cuticle surface microbiota was present in the F_1_ larvae, this heterogeneity was only evident in the cuticle surface microbiota of adult progeny from the same generation. This work provides comprehensive fundamental data for studying how parentage and environmental conditions differentially or concomitantly affect mosquito microbiome composition.

Previous studies on other mosquito species have also shown a loss of field-acquired internal mosquito microbiota following several generations of laboratory colonization [37, 38]. Until now, however, the laboratory generation at which this loss occurs was largely undescribed. Our study represents an initial step in filling this knowledge gap and provides further details regarding the dynamics of changes in both the internal and cuticle surface microbial niches. The homogeneity in internal microbiota of F_1_ adults is suggestive of a loss of field-acquired microbiota, which may have implications for studies that rely on laboratory progeny in lieu of field populations [25, 27, 28]. Previous findings showed that rearing mosquitoes in water from the field could preserve the field-derived internal microbiota in laboratory-colonized adult progeny of *An. gambiae s. l*. for several generations [37]. In addition, other studies, have shown that the microbiota in larval habitat water significantly influences the internal microbiota of emerging adult mosquitoes [40, 41]. Thus, preserving wild-type microbiota in F_1_ progeny could be an avenue for studying mosquito-microbe dynamics in the field.

Maternal egg-smearing has been proposed as a mechanism through which adult female mosquitoes and other insects transfer microbes to their progeny [42-45]. This egg-smearing could explain the heterogeneity observed in internal and cuticle surface microbiota between F_1_ larvae with different maternal origins. On the other hand, a lack of this heterogeneity in the internal microbiota of adult progeny could be attributed to the physiological changes that occur during metamorphosis and adult eclosion, whereby elimination of the larval meconial peritrophic membrane and meconium (midgut and midgut content), along with ingestion of exuvial fluid— which is said to be bactericidal—results in sterile or nearly sterile midguts in newly emerged adults [39, 46]. This could additionally be explained by mechanisms that regulate the composition of adult mosquito internal microbiota [47-49]. Conversely, the heterogeneity in cuticle surface microbiota between adult progeny originating from different maternal sites, suggests that maternally derived microbes in the rearing trays may have colonized adult cuticle surfaces during emergence. The mechanisms underlying the assemblage of the mosquito cuticle surface microbiome are largely undescribed, and thus require further investigation.

The low inter-sample variation in microbial diversity observed in this study has largely been described in laboratory mosquito colonies [38, 50]. The microbial composition of laboratory-reared larvae is typically less diverse [50, 51] compared to those of field-derived larvae, but our laboratory-reared larvae exhibited a rich microbial composition that was comparable to those of field populations [39, 40]. In contrast, our adult progeny had a less diverse microbial composition that was reflective of typical laboratory-reared adult mosquitoes [36, 52]. Further suggesting that field-acquired microbiota, although transferred to laboratory progeny, may be lost within one generation of laboratory colonization—particularly at the adult stage.

In this study, we detected microbial taxa that have previously been identified in *Anopheles* and other mosquito genera [7, 41, 53, 54]. While a majority of the taxa in F_1_ larvae were shared between both the internal and cuticle surface microbial niches, a greater abundance of microbial taxa was detected in the internal microbial niche compared to the cuticle surface. The cuticle surface microbiota of mosquitoes and other hematophagous insects are largely uncharacterized and the mechanisms underlying their assemblage remain unknown. As such, we hypothesize that although both internal and cuticle surface niches are exposed to the same water from which the microbiota is derived, a more conducive and/or selective internal environment could allow for greater proliferation of colonizing bacteria. In F_1_ adults however, less than half of the detected microbial taxa were shared between the internal and cuticle surface microbial niches, suggesting differences in physiological conditions that favor microbial colonization, and corroborating findings that point toward microbial regulatory mechanisms within the mosquito midgut [47-49]. Although a few microbial taxa overlapped between adult internal and cuticle surface microbial niches and the most abundant taxa were shared, many of the unshared taxa have been previously detected in adult mosquitoes including *Anopheles* [1, 54, 55], indicating that the cuticle surface microbiota characterized in this study are inherently associated with mosquitoes. Like the larval microbiota, there was a higher abundance of microbial taxa in the adult internal microbial niche compared to the cuticle surface, further supporting the hypothesis of a more conducive and/or selective internal environment.

With the exception of the adult cuticle surface microbial niche, a majority of all detected microbial taxa overlapped between collection sites in both F_1_ larvae and adults, albeit with differing abundances. This reflects restrictions imposed by controlled laboratory environments in the development of mosquito microbiota. In both microbial niches of both larvae and adults, microbial taxa that were specific to collection sites were low in abundance, compared to the moderate to high abundance of those that were shared across all locations. This was particularly true for *Asaia*—notorius for rapidly colonizing laboratory mosquitoes [56]—which constituted at least 70% of both adult internal and cuticle surface microbiota from progeny across all collection sites. These results suggest that field-acquired mosquito microbiota may be lost in as early as the first generation of laboratory colonization.

We recognize that not having the microbial community profiles of the mothers from which the F_1_ progeny were derived is a limitation of this study. However, the findings herein provide empirical data on the composition of laboratory reared F_1_ *An albimanus* microbiota from different locations. It provides a foundation for exploring the role of parentage, environmental conditions, and inherent host physiological characteristics on the assemblage of the mosquito microbiome, as well as the fate of field-derived microbes upon laboratory colonization. This is critical for advancing mosquito microbiome studies and their applications beyond laboratory settings.

## Methods

The findings presented here extend those of a larger study [8]. Thus, the mosquito collection, processing and sequencing procedures have previously been described in detail [8].

### Mosquito collections and laboratory generation of F_1_ progeny

Gravid and/or blood-fed adult female *An. albimanus* were sampled across four field sites in the villages of Las Cruces and El Terrero, in La Gomera, department of Escuintla, Guatemala (Figure 1). Field-collected mosquitoes from each location were held in separate paper cups, sustained on 10% sucrose solution, and transported to the insectary at Universidad del Valle de Guatemala in Guatemala City for species identification, oviposition and subsequent rearing of F_1_ progeny. Mosquitoes that were morphologically identified as *An. albimanus* following identification keys [57] (approximately 300 in total) were subjected to oviposition, and a subsample of the resulting F_1_ progeny was used for molecular verification of species identity as described below. A modified oviposition procedure [58] was employed, wherein no more than 70 gravid females from the same location were placed in quart size paper ice cream containers with distilled water to a depth of at least 2 cm. The containers were covered with fine mesh fabric that were secured with rubber bands prior to the introduction of gravid field-caught adult *An. albimanus*. The mesh was topped with cotton balls soaked in 10% sucrose to sustain the mosquitoes, and subsequently covered with a thick piece of black plastic bag to keep the containers dark and trap in moisture. The oviposition chambers were held under the following insectary conditions; 27± 2° C, 80± 10% relative humidity, and 12-h light-dark cycle. After at least 48 h, the adult females were removed and discarded as they were not needed for the original study. The F_1_ eggs were collected, pooled by location and reared separately under identical ambient conditions (as described above) and the following larval feeding regimen. Eggs were washed into 18 × 14 × 3 inch plastic larval trays (approximately 200 eggs per tray) containing distilled water to a depth of at least 2 cm, and 3-4 drops of 10% yeast solution. Hatched larvae were sustained on finely ground Koi fish food (Foster & Smith, Inc. Rhinelander, WI) until pupation. At the third to fourth larval instar stage (L3-L4), half of the larvae were separated and used for bioassays in the original study and subsequently processed for microbiota characterization (n=132). Using a stereo microscope, female pupae from the remaining mosquitoes were separated into 8 oz. paper ice cream cups and placed into cardboard cages for adult eclosion. The resulting F_1_ adult virgin female mosquitoes were sustained on 10% sucrose solution until they were 2-5 days old and also used for bioassays in the parent study. These were subsequently processed for microbiota characterization (n=135). Post bioassays, larval and adult samples were preserved in RNALater® solution (Applied Biosystems, Foster City, CA), shipped on dry ice to the US Centers for Disease Control and Prevention (CDC) in Atlanta, USA, and stored at -80 °C until further processing. Samples originating from different geographic locations were handled, stored and processed separately.

### Genomic DNA extraction from mosquito legs, whole mosquito samples, and mosquito cuticle surfaces

Stored mosquito samples were thawed overnight at 4° C, and the thawed RNALater® solution was discarded. This was followed by a rinse with nuclease-free water to remove any residual RNALater®. Legs from a subsample of individual F_1_ adults were removed using sterile forceps and placed in individual sterile1.5 mL Eppendorf tubes containing 75 μL Extracta DNA Prep solution (Quantabio, Beverly, MA) for DNA extraction following manufacturer’s instructions. Both larvae and adult (with and without legs removed) F_1_ samples were pooled for whole body and cuticle surface DNA extractions. Each pool comprised 3 individuals, resulting in 44 pools of larvae and 45 pools of adults. Table 1 shows the number of pools (replicates) processed per life stage and collection site. To obtain cuticle surface samples, each pool was submerged in 500 μL of nuclease free water and washed vigorously by agitating with a vortex mixer for at least 15 seconds. The resulting wash water was transferred to new sterile tubes for DNA extraction. We acknowledge that some microbes may have been lost via rinsing off the preservative (RNALater® solution) prior to this step. A comprehensive (comparable or more than the internal community) microbial community was nonetheless recovered from the cuticle surface samples. The washed sample pools were further surface sterilized using two vigorous washes; first in 70% ethanol, then nuclease free water, each with at least 15 seconds agitation on a vortex mixer. This was followed by one gentle rinse with nuclease free water. Genomic DNA from the surface-sterilized samples and wash water (subsequently referred to as internal and cuticle surface, respectively) was isolated using the TissueLyser II and DNeasy Blood and Tissue Kit (QIAGEN, Hilden, Germany) as follows: in individual 2 mL sterile tubes, 180 μL of buffer ATL (QIAGEN) and 5 mm diameter stainless steel beads (QIAGEN) were added to each sample pool. With the following settings: 30 hz/s for 8 and 15 minutes for internal and cuticle surface samples respectively, samples were homogenized using the TissueLyser (QIAGEN) fitted with two sets of 96-well adapter plates that held the tubes. The plates were rotated every minute during homogenization, and afterwards, samples were transferred into new sterile 1.5 mL Eppendorf tubes for the remaining part of the DNA extraction process. 20 μL of Proteinase K (QIAGEN) was added to each homogenized sample and incubated overnight at 56 ° C, after which 200 μL of buffer AL (QIAGEN) was added and incubated for a further 2 h at the same temperature. The remaining steps were performed according to QIAGEN’s spin-column protocol for purification of DNA from animal tissues, and the purified DNA was eluted in 70 μL of buffer AE (QIAGEN). Two sets of two blank controls (without samples) were processed alongside the internal and cuticle surface samples, and all steps, along with those described below, were performed under sterile conditions. The purified genomic DNA from samples and blank controls were stored at -80 ° C until further processing.

### Molecular species confirmation of *An. albimanus* and 16S rRNA amplicon sequencing

DNA samples from mosquito legs were used as templates to amplify the second internal transcribed spacer region (ITS2) of the mosquito ribosomal DNA in order to verify morphological identification of *An. albimanus*. This was achieved by conventional PCR using the universal ITS2 primers (ITS2 A: TGTGAACTGCAGGACACAT and ITS2 B: TATGCTTAAATTCAGGGGGT) for distinguishing members of *Anopheles* complexes [59]. With a final reaction volume of 25 μL, each PCR comprised ≥ 100 ng/μL DNA template, 15 μ M of each primer, 12.5 μL of 2X AccuStart II PCR SuperMix (Quantabio, Beverly, MA), and PCR grade water to final volume. With the following conditions, initial denaturation at 94 □°C for 4□min, then 35 cycles of 94□°C for 30□s, 53□°C for 40□s, and 72□°C for 30□s, followed by a final extension for 10□min at 72□°C, the reactions were performed using a T100™ Thermal Cycler (Bio-Rad, USA). Using EtBr-stained agarose gel electrophoresis, the amplified products sized ∼500 bp, confirmed all samples as *An. albimanus* [60].

DNA from internal and cuticle surface samples, along with those from blank controls, were used as templates to amplify the universal bacterial and archaeal 16S rRNA gene. This was also achieved by conventional PCR using universal primers targeting the v3-v4 region of the 16S rRNA gene (341F: **TCGTCGGCAGCGTCAGATGTGTATAAGAGACAG**CCTACGGGNGGCWGCAG, and 805R: **GTCTCGTGGGCTCGGAGATGTGTATAAGAGACAG**GACTACHVGGGTATCTAATCC), each with Illumina^®^ (San Diego, CA USA) adapters (bold typeface). With a final reaction volume of 25 μL, each PCR comprised; ≥20□ng/µL DNA template, 5□µM of each primer, 10□µL of 2× KAPA HiFi HotStart PCR mix (Roche, Switzerland), and PCR grade water to final volume. Three negative controls with PCR grade water substituted for DNA template, were processed along with the samples. The reactions were performed using the T100™ Thermal Cycler (Bio-Rad, USA) under the following conditions; initial denaturation at 95 °C for 3□min, then 25 cycles of 95□°C, 55□°C, and 72□°C for 30□s each, followed by a final extension for 5□min at 72□°C. The amplification products, sized ∼460 bp, were verified by electrophoreses as described above and quantified using a NanoDrop™ spectrophotometer (Thermo Fisher Scientific, Waltham, MA). Amplicons, including PCR products from blanks and negative controls, which yielded no bands following electrophoresis, were purified using Agencourt AMPure XP beads (Beckman Coulter Inc., Indianapolis, IN, USA) at 0.7X (internal) or 0.875X (cuticle surface) sample volume, and eluted in 40□µL of 10□mM Tris buffer (pH 8.5).

All purified products, including those of blanks and negative controls, were used as templates for sequencing library preparation. This was accomplished via index PCR using the Nextera XT Index kit v2 sets, A, B and D (Illumina, San Diego, CA). With a final reaction volume of 50 μL, each index PCR comprised; 25□µL NEBNext High-Fidelity 2X PCR master mix (New England Biolabs Inc., Ipswich, MA), 5□µL of each index primer, 10□µL of purified PCR products (0– 20□ng/µL) as template, and PCR grade water to final volume. The PCRs were performed under the following reaction conditions; 98□°C for 30□s, then 8 cycles of 98□°C for 10□s, 55□°C and 65□°C for 30□s each, and a final extension at 65□°C for 5□min, and resulting libraries were also cleaned using Agencourt AMPure XP beads (Beckman Coulter Inc., Indianapolis, IN, USA) at 1.2X sample volume and eluted in 25□µL of 10□mM Tris buffer (pH 8.5). The libraries were subsequently analyzed for size and concentration, normalized to 2□nM, pooled and denatured following Illumina guidelines for loading onto flowcells. Sequencing was performed on an Illumina HiSeq 2500 machine, using 2×250 cycle paired-end sequencing kits.

### Preprocessing of sequencing reads

Sequencing outputs were demultiplexed and converted to the fastq format for downstream analysis using the bcl2fastq (v2.19) conversion software (Illumina®). A total of 115,250,077 demultiplexed paired-end sequencing reads, with a maximum length of 250 bp were initially imported into the ‘quantitative insights into microbial ecology’ pipeline, QIIME2 v.2017.7.0 [61], and further sequencing read processing and analysis were performed in v.2018.2.0 of the pipeline. Using the DADA2 plugin in QIIME 2 [62], the `denoise-paired` command with the following options; `trunc_len_f: 244, trunc_len_r: 244, and n_reads_learn: 500000`, was used to correct errors, remove chimeras and merge paired-end reads. The resulting amplicon sequence variants (ASVs; n=30,956,883) were further filtered to remove potentially extraneous ASVs (those with <100 counts) and ASVs that were associated with blanks and negative controls. This last filtering step resulted in 17,225,776 ASVs, ranging from 3,277 to 223,222 per sample (mean 96,774), that were used for downstream comparison of bacterial composition and taxonomic analysis. Suppl. 3. shows sequencing reads and ASV summary statistics.

### Diversity indices

Analysis of microbial diversity within (alpha diversity) and between (beta diversity) samples were performed in QIIME2 using the Shannon diversity index and Bray-Curtis dissimilarity index, respectively. The Shannon diversity indices were calculated using rarefied ASVs counts per sample, in which ASVs per sample were selected randomly without replacement at an even depth (Suppl. 4) for ten iterations. The resulting average Shannon indices are presented and were compared between samples using pairwise Kruskal-Wallis tests with Benjamini-Hochberg false discovery rate (FDR) corrections for multiple comparisons.

The Bray-Curtis dissimilarity indices were computed with or without rarefaction, and resulting indices were compared between samples using pairwise PERMANOVA tests (999 permutations) with FDR corrections. There were no discernable differences between results of rarefied and non-rarefied data. Thus, results of Bray-Curtis dissimilarity indices using non-rarefied data were visualized by Principal Co-ordinates Analysis (PCoA) plots in R [63] using the phyloseq R package [64].

Significance for both pair-wise analyses was set to *q* <0.05 (i.e. post FDR *p*-value corrections).

### Taxonomic analysis and differentially abundant microbial taxa

Taxonomic analysis of ASVs was performed using QIIME2’s pre-trained Naïve Bayes classifier [65] and q2-feature-classifier plugin [66]. Prior to analysis, the classifier was trained on the QIIME-compatible 16S SILVA reference (99% identity) database v.128 [67], and using the extract-reads command of the `q2-feature-classifier` plugin, the reference sequences were trimmed to the v3-v4 region (425 bp) of the 16S rRNA gene. The relative abundance of annotated ASVs across samples were subsequently visualized using the `qiime feature- table heatmap` plugin based on Bray-Curtis distances with the plugin’s default clustering method. Only annotated ASVs with counts ≥ 2000 (larvae) or ≥ 1000 (adults) were included in the heatmaps.

Differentially abundant microbial taxa across locations were identified using QIIME2’s analysis of composition of microbiomes (ANCOM) [68] plugin. The cut-off for differential abundance was set to an effect size of log F≥20 and W≥20, i.e. a taxon was differentially abundant across collection sites if the ratio of its abundance to those of at least 20 other taxa (25% of all included taxa) differed significantly across sites.

Prior to each analysis, ASV frequency data was normalized by log_10_ transformation following the addition of pseudocounts of 1. To ensure that filtering of low frequency reads did not compromise our findings, all downstream analyses were also performed with these reads included, and results (Suppl 5) were consistent with those of the clean/filtered dataset. Results of the clean dataset are subsequently discussed.

The outputs of data analyses were aesthetically formatted using Inkscape [69].

## Declarations

### Ethics approval and consent to participate

Not applicable

### Consent for publication

Not applicable

### Availability of data and material

The raw sequencing reads generated from this project, including those from negative controls (blanks), have been deposited in the National Center for Biotechnology Information (NCBI), Sequence Read Archive under the BioProject PRJNA512122.

### Author contributions

ND^†^ & AL conceptualized and designed the study; NP facilitated and provided facilities for field work; ND^†^, ACB, FL & JCL performed mosquito collections, and mass rearing; ND^†^ & MS performed molecular analysis and sequencing; ND^†^ analyzed the data; ND^†^ and ND performed the data visualizations; ND^†^ drafted the manuscript; all authors reviewed and approved the final version of the manuscript.

## Acknowledgements

We thank the cattle coral owners in Escuintla, Guatemala for permission to conduct mosquito collections on their corals, without which this study would not have been possible; the Malaria Research and Reference Reagent Resource Center (MR4) for providing the ITS2 primers used for *Anopheles albimanus* species identification; Nelson Jimenez and Ricardo Santos from the Ministerio de Salud Publica y Asistencia Social (MSPAS) for assistance during mosquito collections; Daniela Da’Costa, Pedro Peralta, Adel Mejia and Alfonso Salam from Universidad del Valle de Guatemala (UVG) for field support and assistance during mosquito rearing; and appreciate comprehensive feedback from two anonymous reviewers on an earlier version of this manuscript—these have (re)shaped how the findings herein are presented and discussed. The adult mosquito icon used in Fig. 1 was created by MarkieAnn Packer from the Noun Project.

The findings and conclusions in this paper are those of the authors and do not necessarily represent the official position of the US Centers for Disease Control and Prevention (CDC) or the American Society for Microbiology (ASM).

## Supporting Information

**Suppl. 1**. Shannon alpha diversity plots

**Suppl. 2**. Frequency and relative abundance of microbial taxa detected in internal and cuticle surface microbiota of F_1_ laboratory colonized *An. albimanus* larvae and adults

**Suppl. 3**. Summary statistics of sequencing reads and amplicon sequence variants (ASVs) used for downstream analysis

**Suppl. 4**. Rarefaction depth and plots of sequencing reads

**Suppl 5**. Results of downstream analysis including low frequency (potentially extraneous) reads

