## Supplementary material for "Comprehensive characterization of internal and cuticle surface microbiota of laboratory-reared F_1_ *Anopheles albimanus* originating from different sites": Suppl. 1

### Slide 1
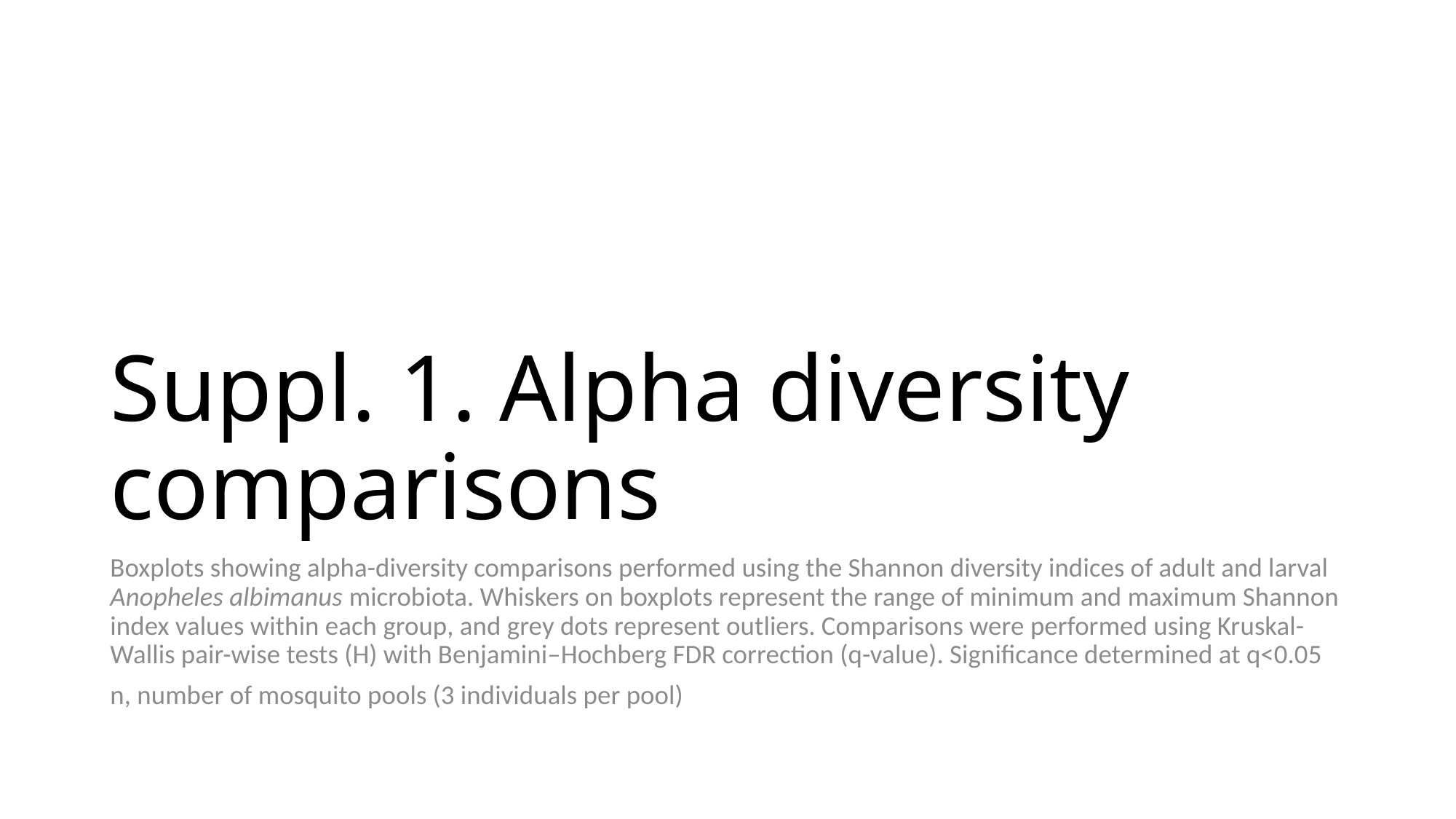

#### Suppl. 1. Alpha diversity comparisons
Boxplots showing alpha-diversity comparisons performed using the Shannon diversity indices of adult and larval Anopheles albimanus microbiota. Whiskers on boxplots represent the range of minimum and maximum Shannon index values within each group, and grey dots represent outliers. Comparisons were performed using Kruskal-Wallis pair-wise tests (H) with Benjamini–Hochberg FDR correction (q-value). Significance determined at q<0.05
n, number of mosquito pools (3 individuals per pool)

### Slide 2
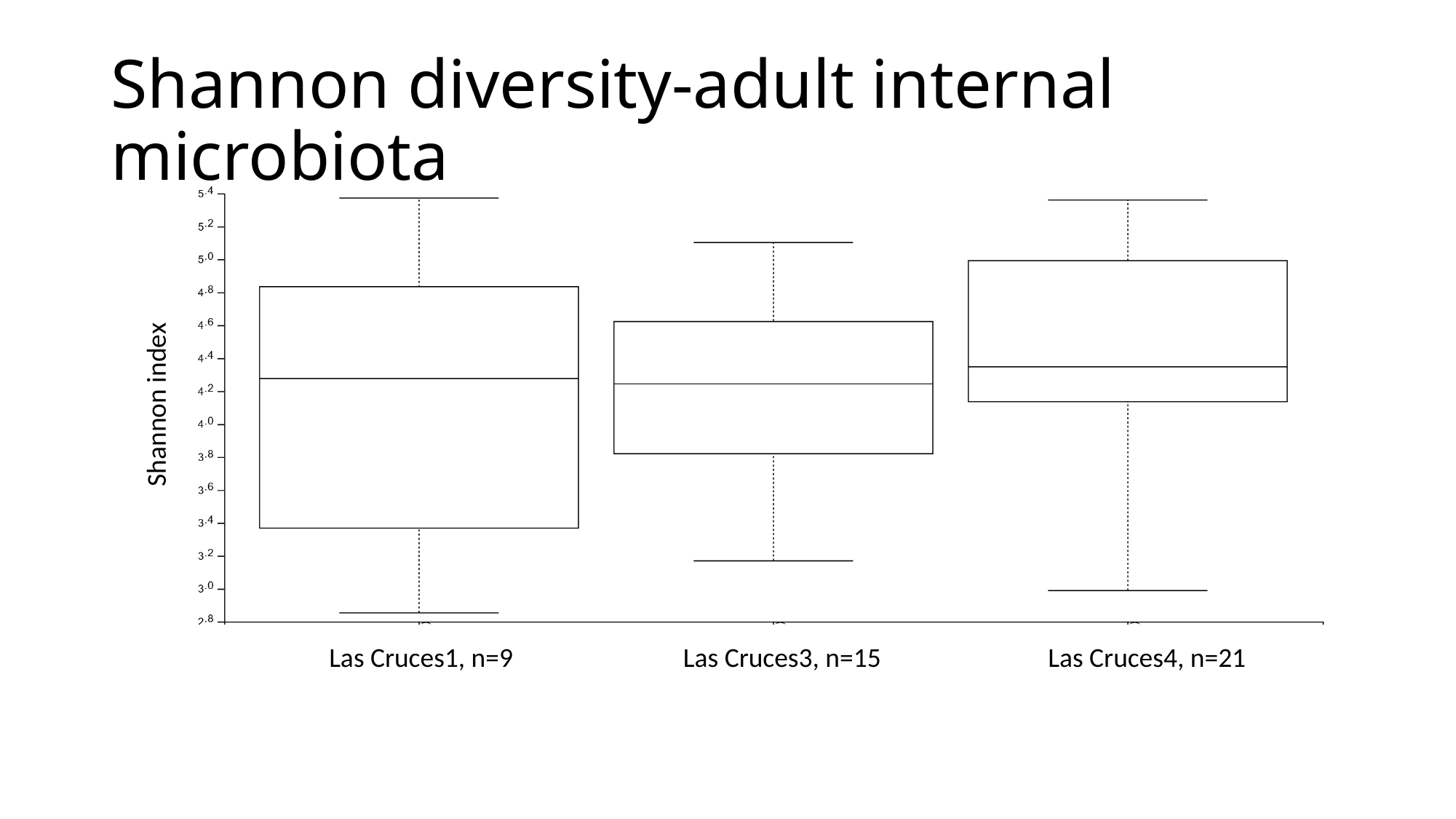

#### Shannon diversity-adult internal microbiota
Shannon index
Las Cruces1, n=9
Las Cruces3, n=15
Las Cruces4, n=21

### Slide 3
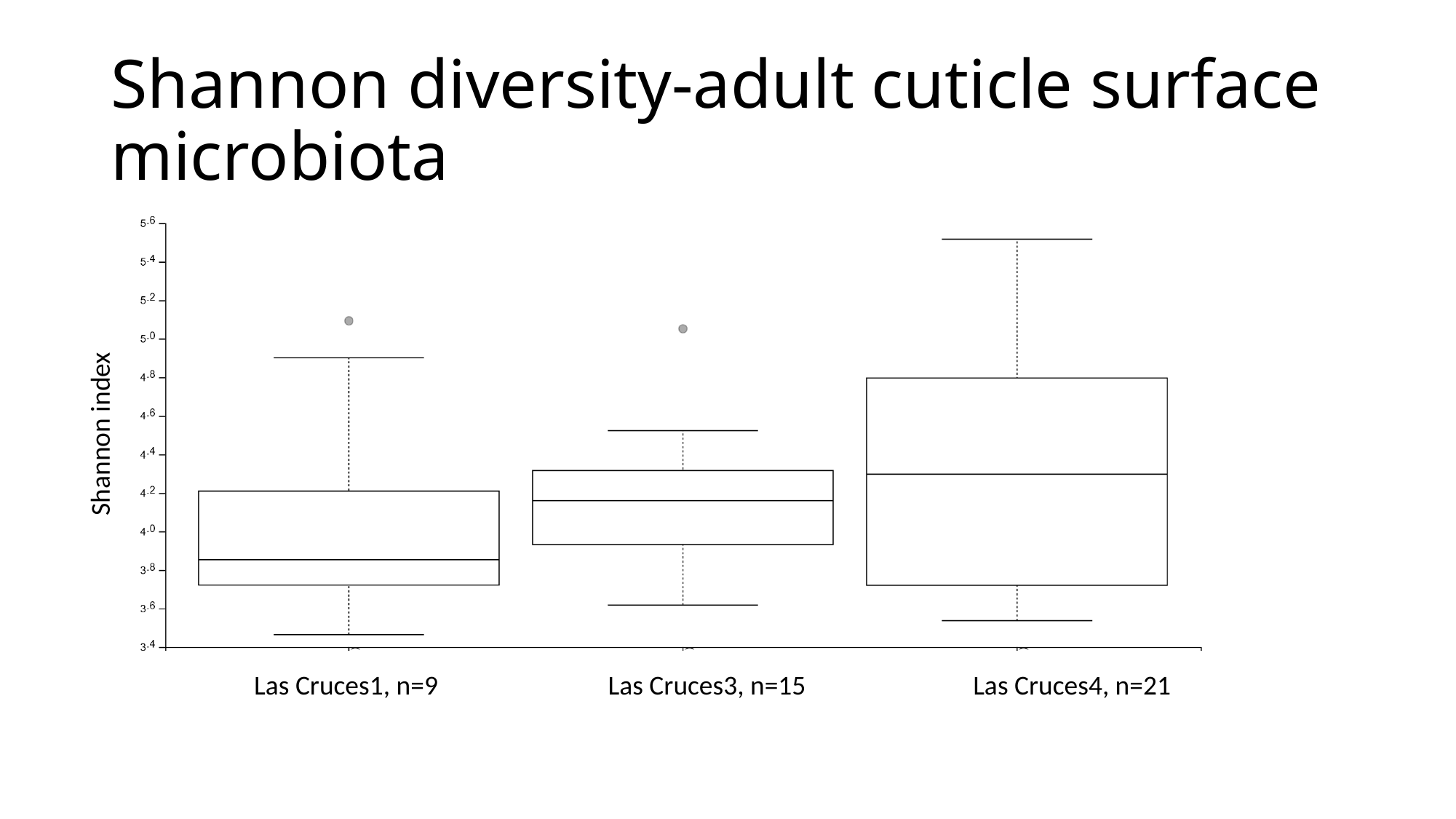

#### Shannon diversity-adult cuticle surface microbiota
Shannon index
Las Cruces1, n=9
Las Cruces3, n=15
Las Cruces4, n=21

### Slide 4
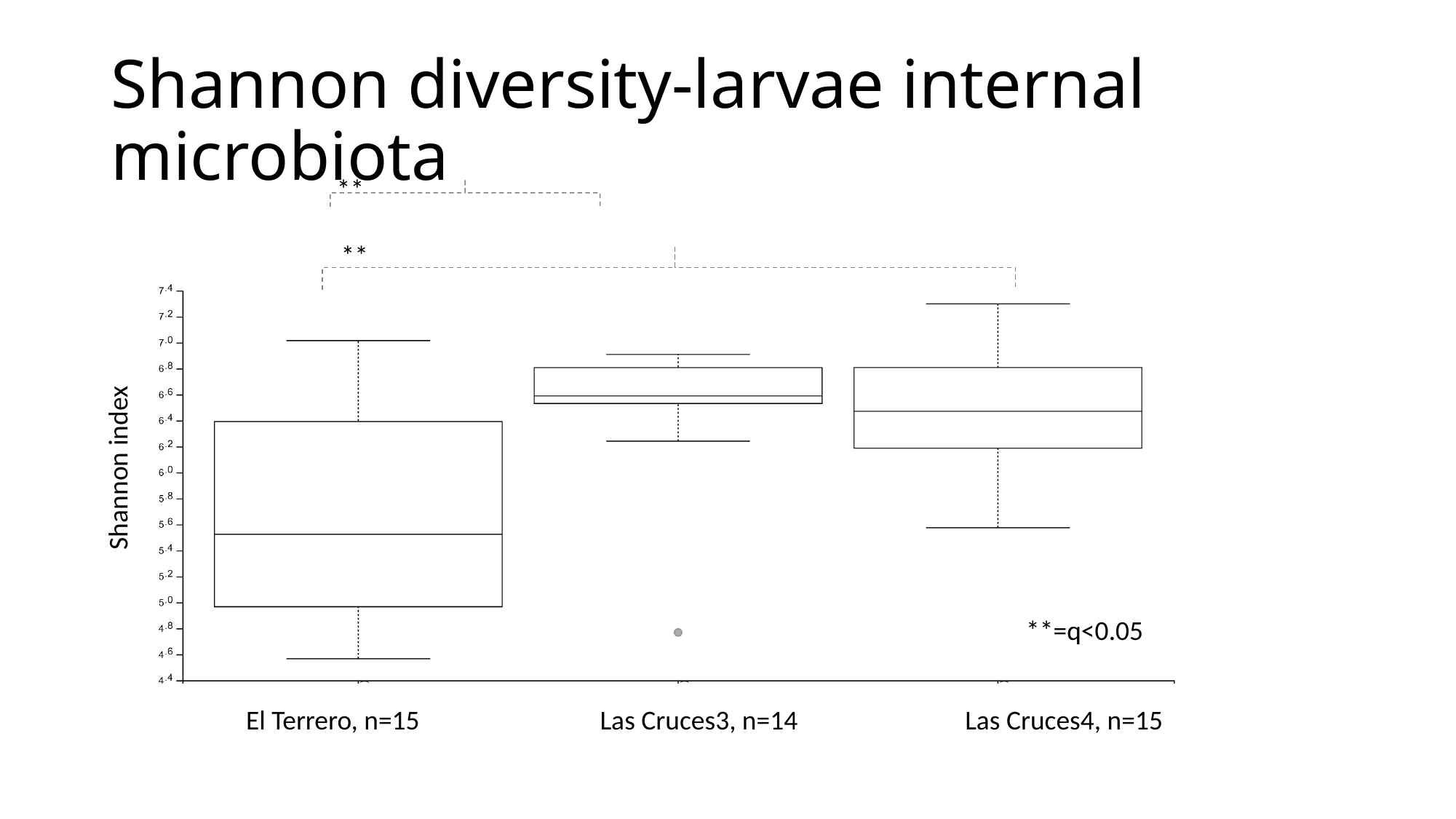

#### Shannon diversity-larvae internal microbiota
**
**
Shannon index
**=q<0.05
El Terrero, n=15
Las Cruces3, n=14
Las Cruces4, n=15

### Slide 5
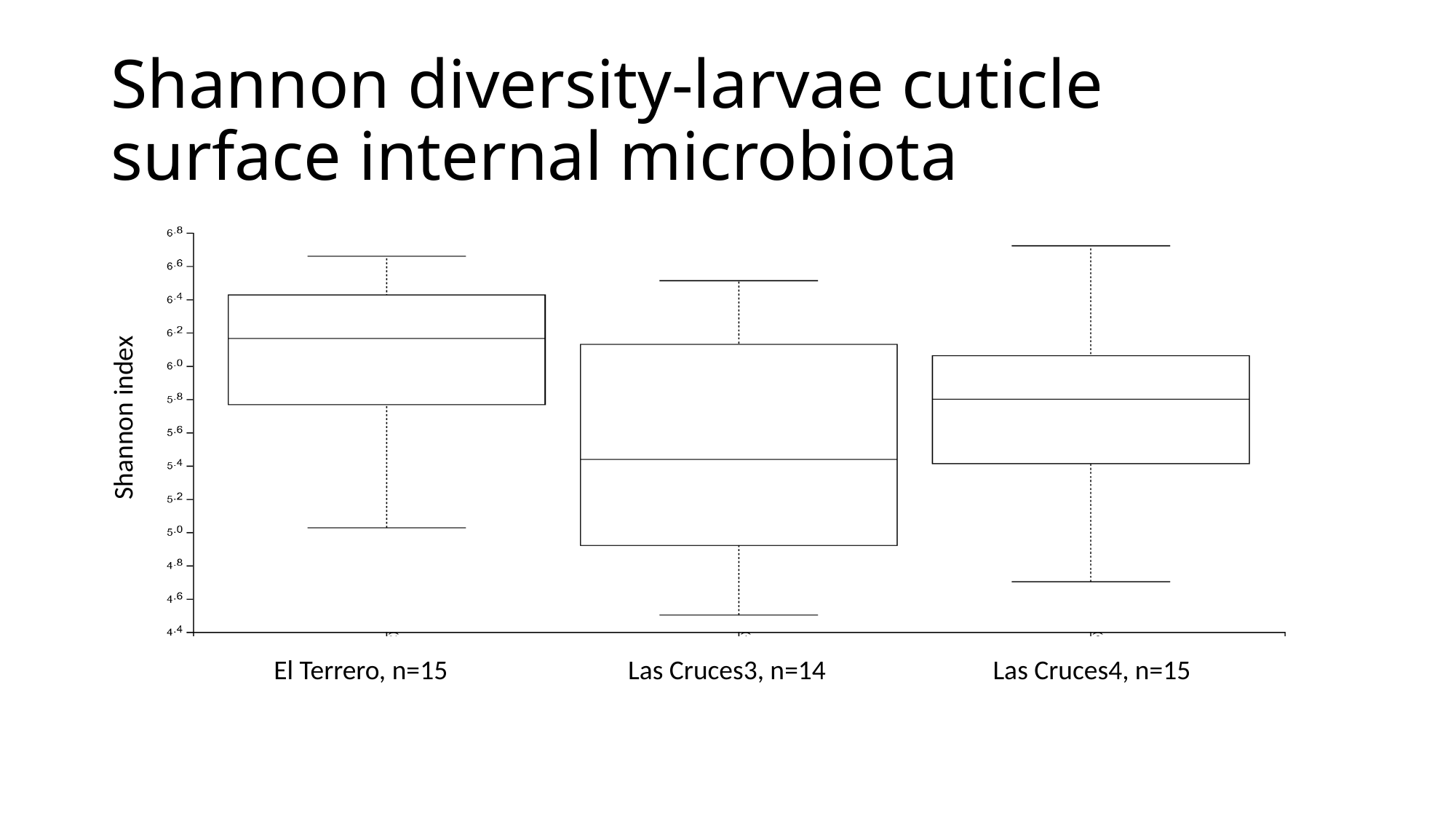

#### Shannon diversity-larvae cuticle surface internal microbiota
Shannon index
El Terrero, n=15
Las Cruces3, n=14
Las Cruces4, n=15
