## Supplementary material for "Comprehensive characterization of internal and cuticle surface microbiota of laboratory-reared F_1_ *Anopheles albimanus* originating from different sites": Suppl. 3

**Suppl. 3: Summary statistics of sequencing reads and amplicon sequence variants (ASVs) used for downstream analysis.**

**a.**

|  | Larvae | |  | Adults | |
| --- | --- | --- | --- | --- | --- |
| **Location** | Internal | Cuticle surface |  | Internal | Cuticle surface |
| El Terrero | 12771326 | 5811542 |  | - | - |
| Las Cruces 1 | - | - |  | 7273037 | 5041136 |
| Las Cruces 3 | 11787864 | 8901204 |  | 12022485 | 8151470 |
| Las Cruces 4 | 12150961 | 8592908 |  | 12864326 | 9848007 |
| **Total** | **36710151** | **23305654** |  | **32159848** | **23040613** |

**b.**

|  | Larvae | |  | Adults | |
| --- | --- | --- | --- | --- | --- |
| **Location** | Internal | Cuticle surface |  | Internal | Cuticle surface |
| El Terrero | 2040105 | 1250090 |  | - | - |
| Las Cruces 1 | - | - |  | 2170546 | 1613411 |
| Las Cruces 3 | 1859321 | 2094032 |  | 4055241 | 3459254 |
| Las Cruces 4 | 1839621 | 2589365 |  | 4213701 | 3767459 |
| **Total** | **5739047** | **5933487** |  | **10439488** | **8840124** |

**c.**

|  | Larvae | |  | Adults | |
| --- | --- | --- | --- | --- | --- |
| **Location** | Internal | Cuticle surface |  | Internal | Cuticle surface |
| El Terrero | 1546552 | 1072094 |  | - | - |
| Las Cruces 1 | - | - |  | 1113910 | 699266 |
| Las Cruces 3 | 1556738 | 1932883 |  | 1678062 | 939002 |
| Las Cruces 4 | 1412494 | 2286499 |  | 1806357 | 1181919 |
| **Total** | **4515784** | **5291476** |  | **4598329** | **2820187** |

Number of sequencing reads and ASVs from the internal and cuticle surface microbiota of whole larvae (n=132) and adult (135) F_1_ progeny of wild caught *Anopheles albimanus* from four locations in La Gomera, Guatemala. Mosquitoes were consolidated into 44 and 45 pools (3 mosquitoes per pool) of larvae and adults, respectively. A total of 55,200,461 (adults) and 60,015,805 (larvae) sequencing reads were obtained and processed for downstream analysis. The table provides the numbers of: **a.** sequencing reads, **b.** ASVs post quality control and dereplication, and **c.** ASVs post removal of low frequency (<100) ASVs and ASVs from blanks per maternal collection site, life stage and microbial niche. Blank controls produced 33,811 sequencing reads, resulting in 4,737 ASVs that were subtracted from each sample prior to downstream analysis.
