## Supplementary material for "Comprehensive characterization of internal and cuticle surface microbiota of laboratory-reared F_1_ *Anopheles albimanus* originating from different sites": Suppl. 4

### Slide 1
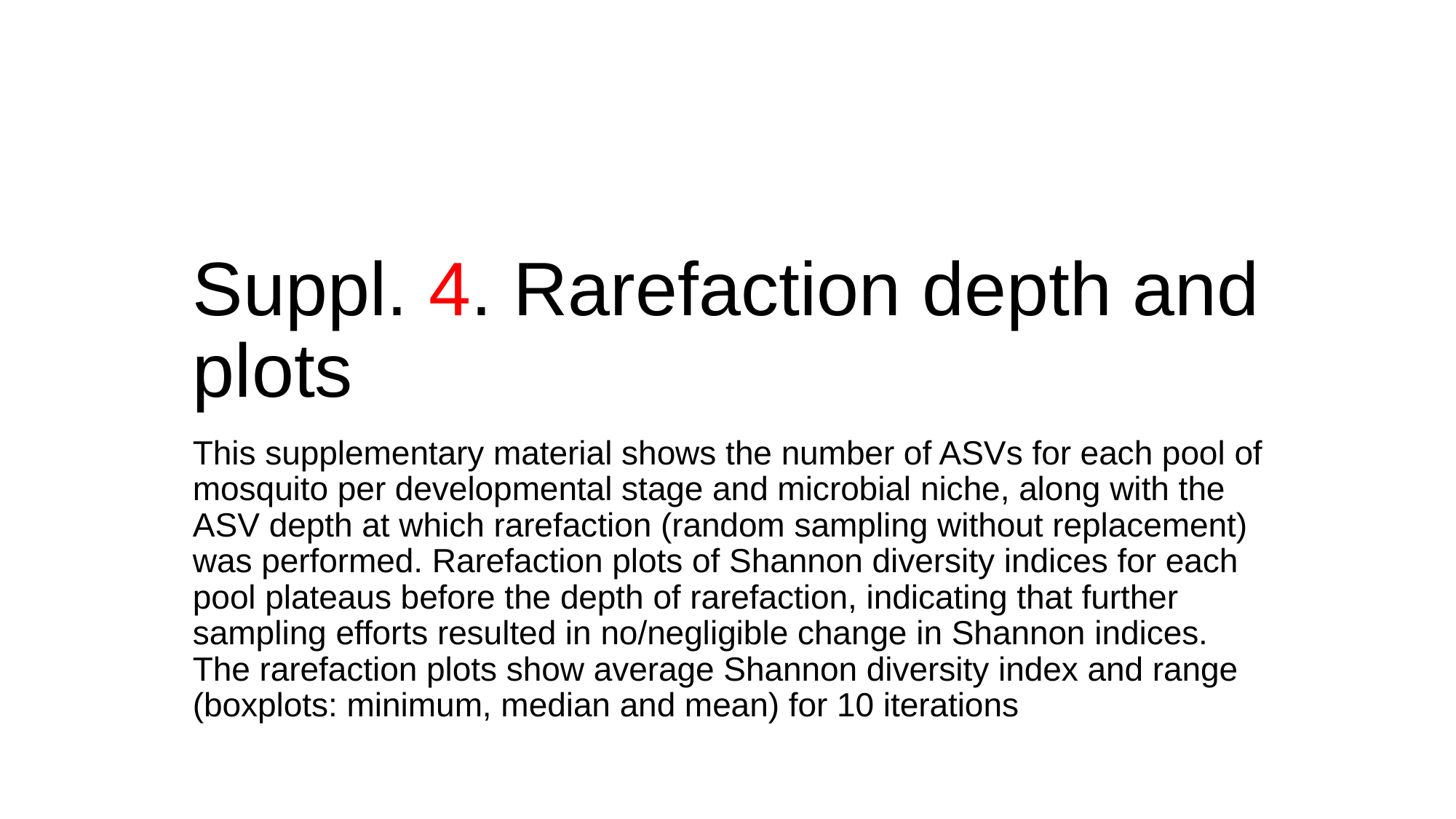

#### Suppl. 4. Rarefaction depth and plots
This supplementary material shows the number of ASVs for each pool of mosquito per developmental stage and microbial niche, along with the ASV depth at which rarefaction (random sampling without replacement) was performed. Rarefaction plots of Shannon diversity indices for each pool plateaus before the depth of rarefaction, indicating that further sampling efforts resulted in no/negligible change in Shannon indices. The rarefaction plots show average Shannon diversity index and range (boxplots: minimum, median and mean) for 10 iterations

### Slide 2
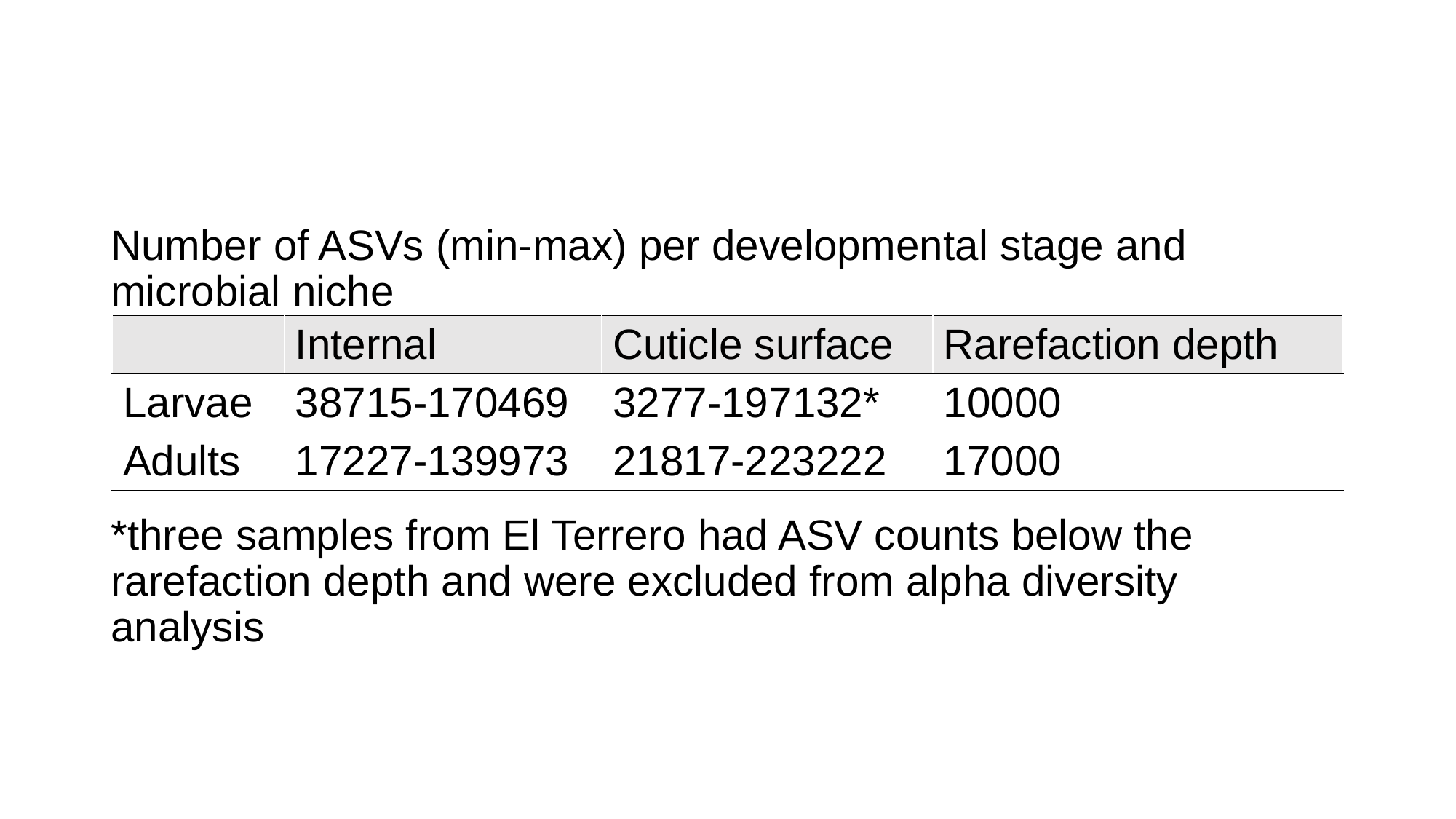

Number of ASVs (min-max) per developmental stage and microbial niche
*three samples from El Terrero had ASV counts below the rarefaction depth and were excluded from alpha diversity analysis
| | Internal | Cuticle surface | Rarefaction depth |
| --- | --- | --- | --- |
| Larvae | 38715-170469 | 3277-197132\* | 10000 |
| Adults | 17227-139973 | 21817-223222 | 17000 |

### Slide 3
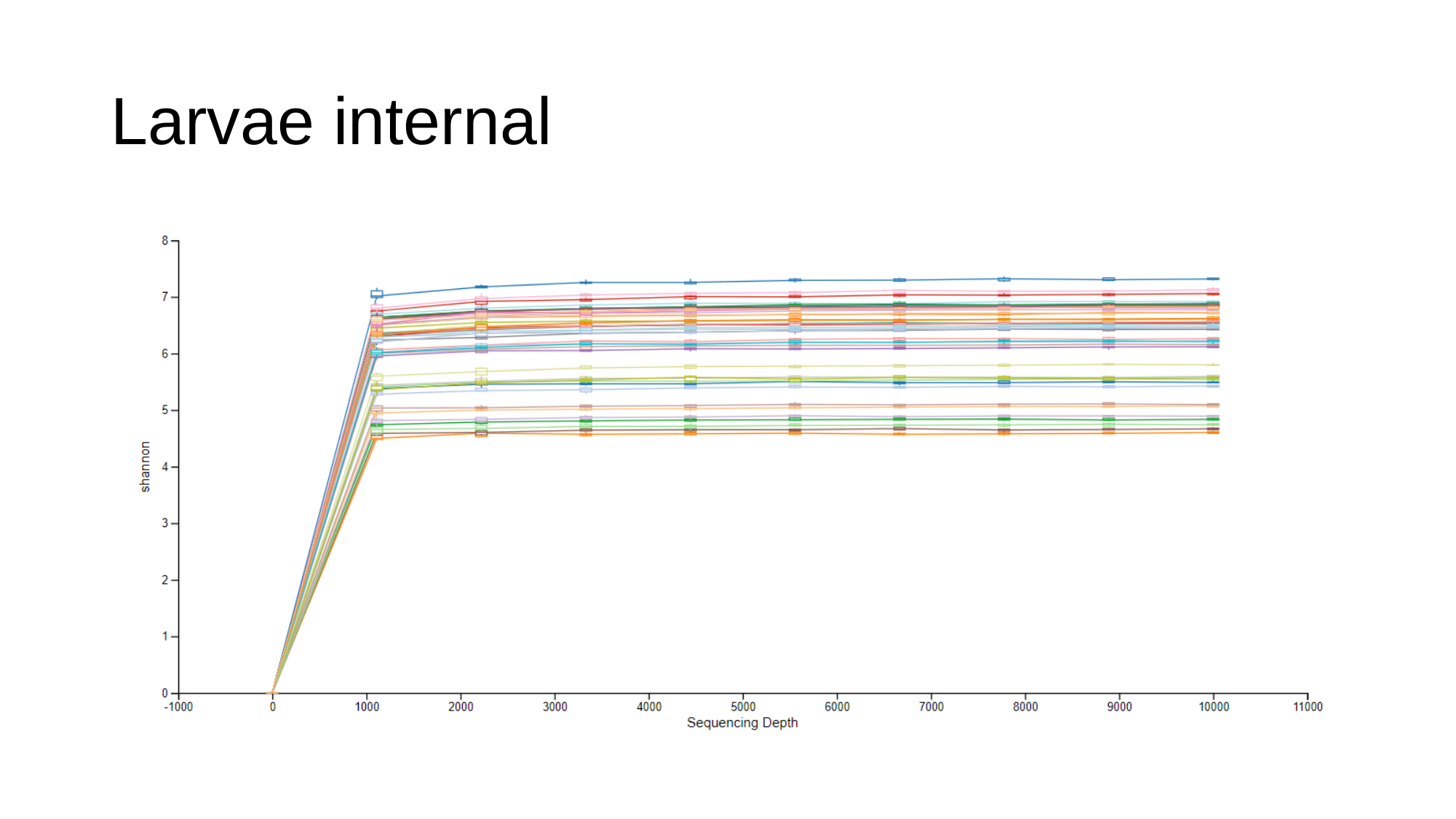

#### Larvae internal

### Slide 4
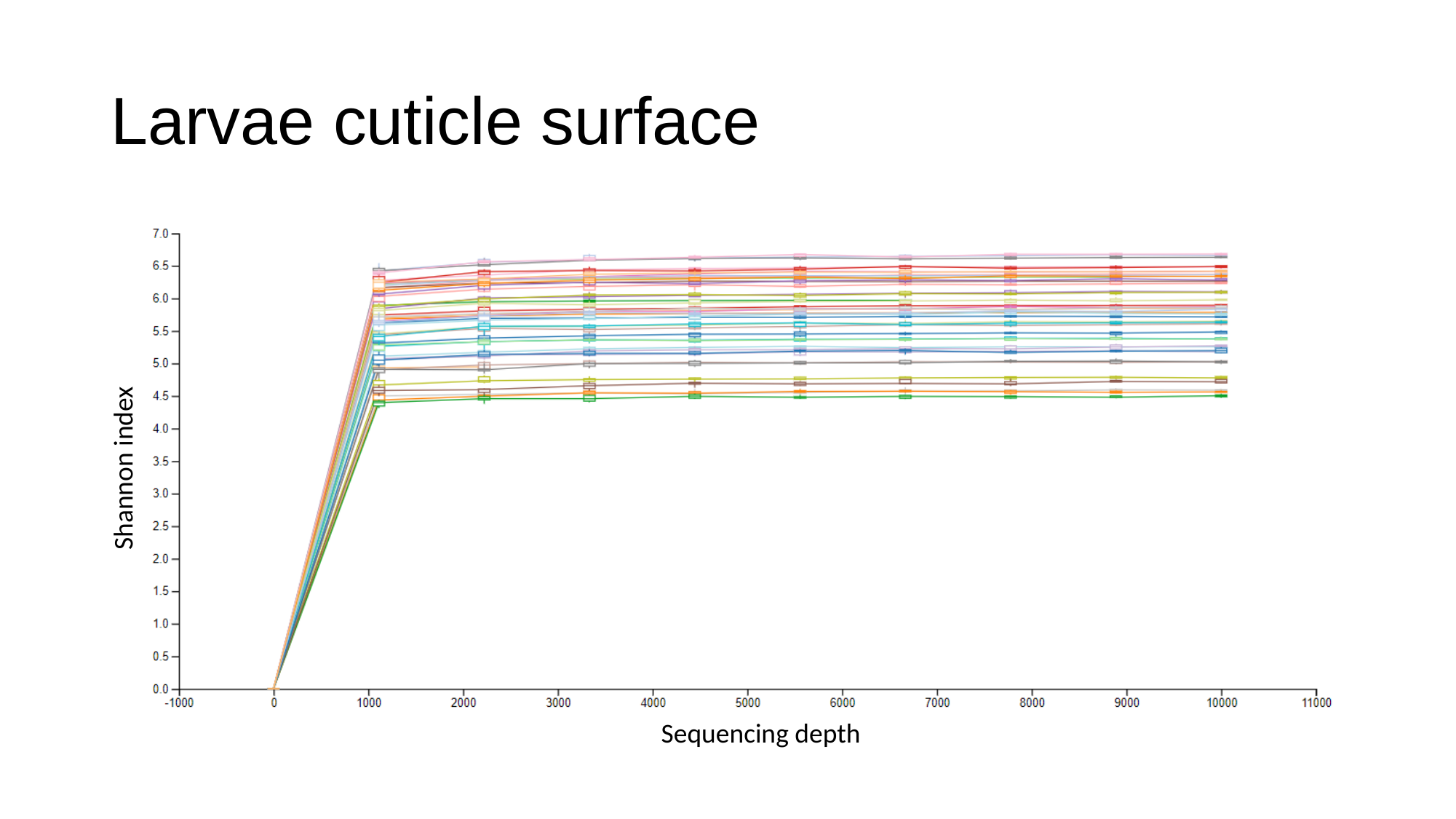

#### Larvae cuticle surface
Shannon index
Sequencing depth

### Slide 5
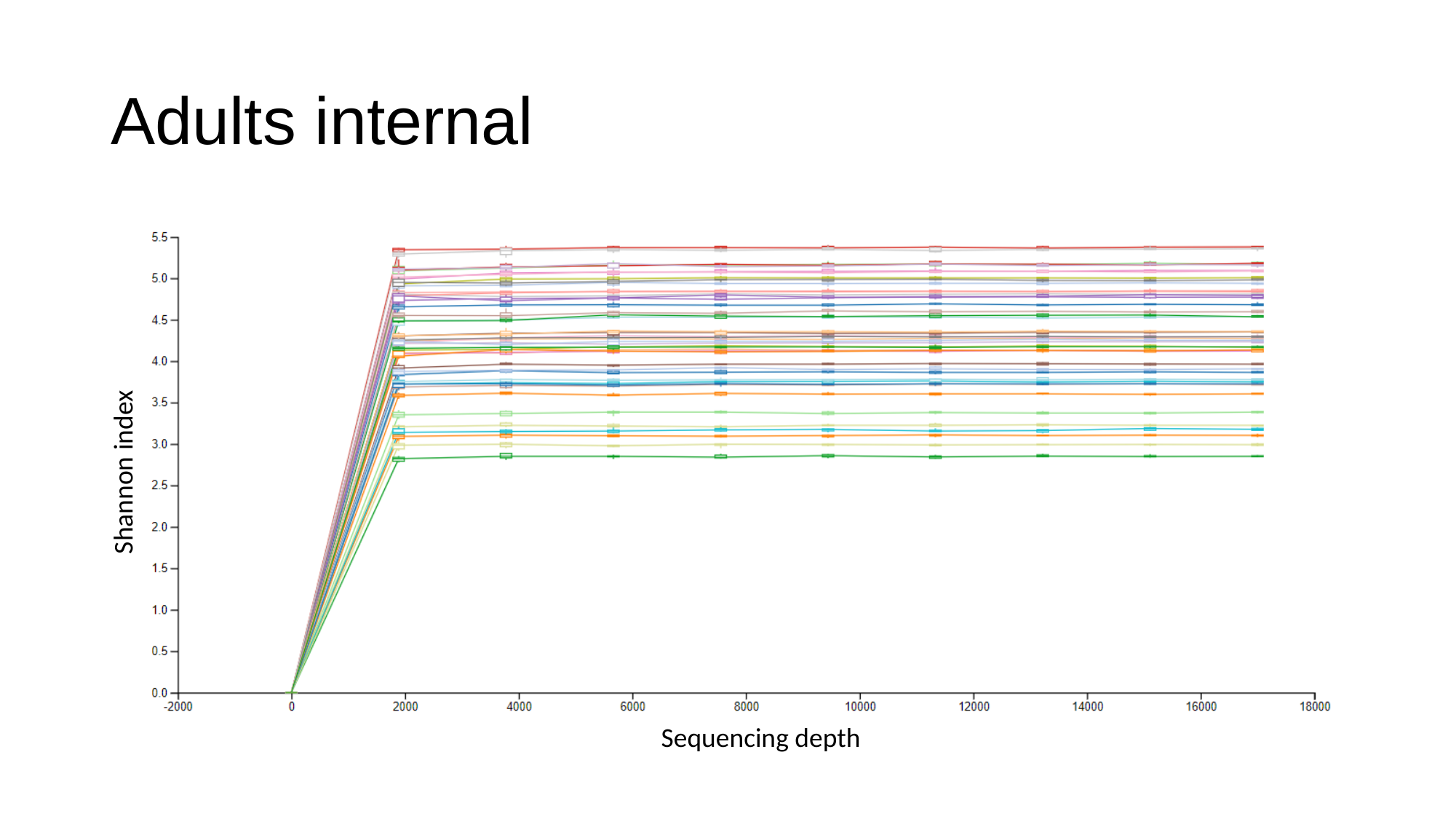

#### Adults internal
Shannon index
Sequencing depth

### Slide 6
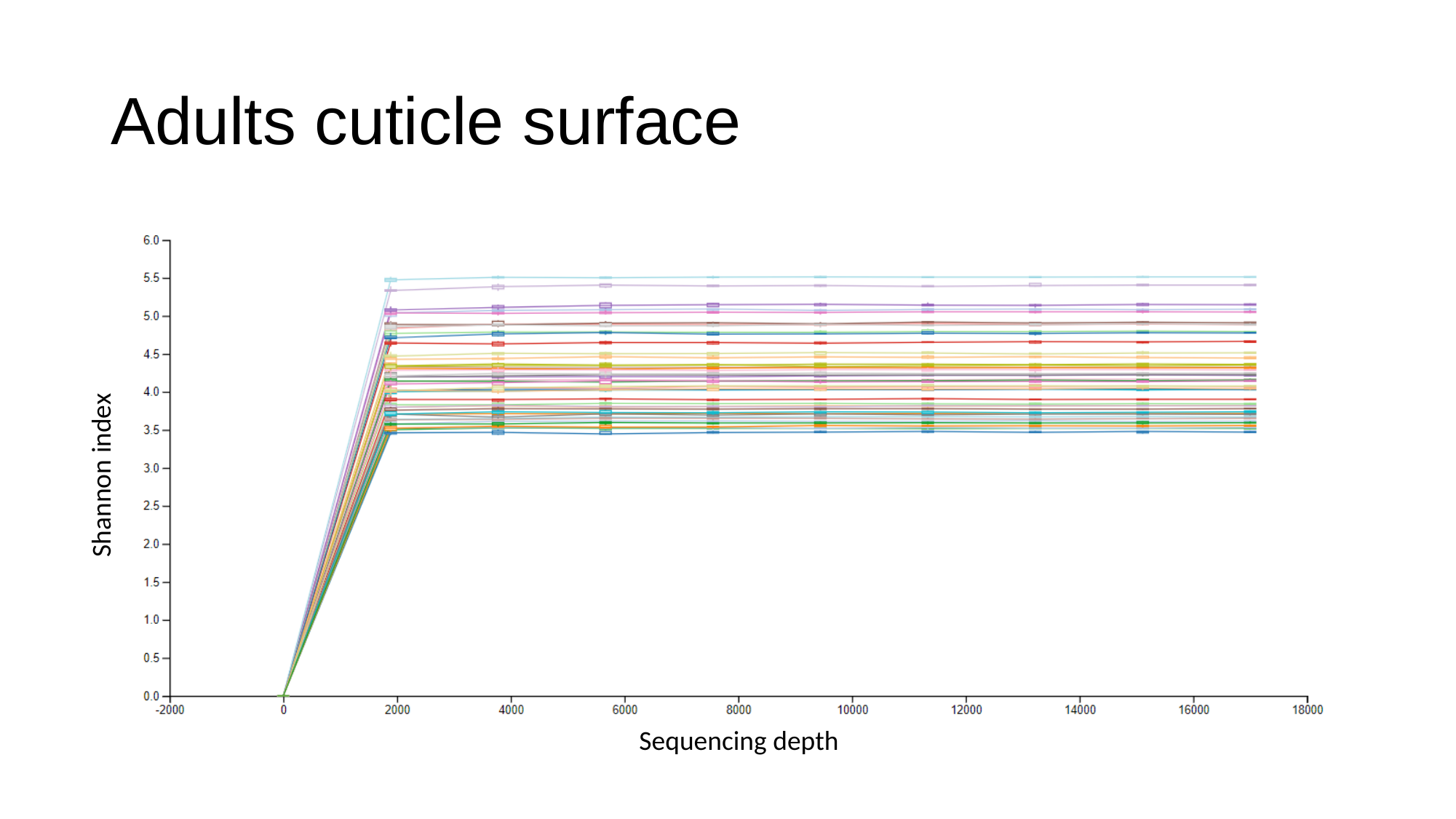

#### Adults cuticle surface
Shannon index
Sequencing depth
